# A Parametric Blueprint for Optimum Cochlear Outer Hair Cell Design

**DOI:** 10.1101/2022.10.11.511831

**Authors:** Richard D. Rabbitt, Tamara C. Bidone

## Abstract

The present work examines the hypothesis that cochlear outer hair cell (OHC) properties vary in precise proportions along the tonotopic map to optimize electro-mechanical power conversion. We tested this hypothesis using a very simple model of a single isolated OHC driving a mechanical load. Results identify three nondimensional ratios that are predicted to optimize power conversion: the ratio of the resistive-capacitive (RC) corner to the characteristic frequency (CF), the ratio of nonlinear to linear capacitance, and the ratio of OHC stiffness to cochlear load stiffness. Optimum efficiency requires all three ratios to be universal constants, independent of CF and species. The same ratios are cardinal control parameters that maximize power output by positioning the OHC operating point on the edge of a dynamic instability. Results support the hypothesis that OHC properties evolved to optimize electro-mechanical power conversion. Identification of the RC corner frequency as a control parameter reveals a powerful mechanism used by medial olivocochlear efferent system to control OHC power output. Results indicate the upper frequency limit of OHC power output is not constrained by the speed of the motor itself, but instead is likely limited by the size of the nucleus and membrane surface area available for ion-channel expression.

## INTRODUCTION

The appearance of outer hair cells (OHCs) in the cochlea roughly 125 million years ago [1] endowed mammals with the ability to hear high frequency sounds, extending > 100kHz in whales and >70kHz in bats [2, 3]. In contrast, hearing in reptiles, amphibians, lizards and fish is typically limited to < 2 kHz [4–6], and hearing in most birds is limited to < 8 kHz [7, 8]. Across vertebrates there is an active process in inner ear sensory hair bundles that plays an important role in sensitivity [9–11], but the key difference in mammals is active electro-mechanical amplification by the cell body of cochlear OHCs [12, 13]. Amplification is enabled by expression of the transmembrane protein prestin [14], and manifested on the whole-cell level as a piezoelectric motor in the lateral wall membrane [15–18]. The structure of prestin suggests motor function likely arises from voltage-driven conformational changes in the protein [19–21], but the detailed molecular mechanism(s) underlying whole cell piezoelectricity remains a subject of research. Prestin is essential for sensitive hearing in mice over the 2-50 kHz range [22, 23], though prestin-independent electromotility has been implicated at frequencies >50kHz [24]. Direct electro-mechanical coupling in OHCs allows electrical power entering the cell primarily through the mechanoelectrical transduction (MET) channels to be converted into mechanical power output, thereby drawing power from the electro-chemical endolymphatic potential to amplify vibrations in the cochlea. Although electrical charge displacement in prestin-expressing membrane patches [25, 26] and voltage-driven whole-cell displacement [27, 28] have low-pass characteristics, both electrical power consumption and mechanical power output are band-pass with best frequency exceeding 50kHz under some conditions [29]. It is currently not known if band-pass electromechanical power conversion in OHCs is tuned to the cochlear tonotopic map, or what role OHC biophysical properties play in setting the best power conversion frequency.

Monotonic variations in OHC biophysical properties along the tonotopic map provide a hint that each OHC might be tuned for best operation at a specific characteristic frequency (CF). Cells located at the 0.2 kHz CF place have a length of ~80 μm while cells located at the 10 kHz CF place have a length of ~40 μm [30–32]. The correlation between OHC length and CF is universal across mammalian species, but precisely why the correlation exists is not known. Whole-cell OHC axial stiffness also varies with cell length [33], and has an inverse log-linear scaling with CF similar to the stiffness of the in-tact cochlear partition [34]. Passive capacitance and peak nonlinear capacitance both scale with cell length, roughly in proportion to the membrane surface area [35, 36], while basolateral ion channel expression and membrane conductance increases with CF [37–39]. Capacitance and conductance combine to cause the passive resistive-capacitive (RC) corner frequency of the cell membrane to increase with CF, which likely plays an important role in amplification by shifting the timing of electro-mechanical force production [40, 41]. These correlations motivate the present hypothesis that specific properties are required for optimum OHC operation at CF. It has been suggested previously that OHC biophysical properties vary with CF to optimize impedance matching [42, 43], or to set the OHC operating point near a dynamic instability [44–46]. Both hypotheses might be true, but the specific optimization problem that nature solved to specify OHC parameters remains unclear.

Here, we present theoretical evidence that OHC biophysical properties maximize electro-mechanical power conversion at CF, achieved by simultaneously matching the impedance of the OHC to the cochlear load *and* by positioning the operating point of the cell on the stable edge of a dynamic instability (the critical point). Results suggest OHCs evolved under rigid constraints relating OHC length and biophysical properties to each other and to the mechanics of the organ of Corti along the tonotopic map. Mathematical equations completely specifying the model are provided in the Methods Section and symbolic derivations are provided in a supplement (Mathematica Code, Wolfram Research). Readers interested in the *critical point conditions* and *impedance matched conditions* that specify optimum OHC design parameters should consult the Results Section. The Discussion Section describes relevance to cochlear amplification and efferent control.

## METHODS

We used the minimal model illustrated in Fig. 1 to examine electro-mechanical power conversion by an isolated piezoelectric OHC working against a passive spring-mass-damper mechanical load. The model was designed to reveal dependence of OHC mechanical and biophysical properties on the load, and is not intended to reproduce tuning in the cochlea. The active process in this model includes only the prestin-dependent lateral wall motor and excludes contributions from hair bundle motility. The coupled system is driven by a force applied to the apical mass, as an analog to sound-induced pressure in the cochlea. Amplification is powered by the mechano-electrical transduction (MET) current entering the cell, which is gated by displacement of the apical end of the cell. The MET current and the piezoelectric coefficient both include saturating nonlinearities. Two different loading conditions are considered to determine if the critical point conditions and/or the impedance matched conditions depend on details of the cochlear load. A complete mathematical description of the model is provided below.

**Fig. 1.**
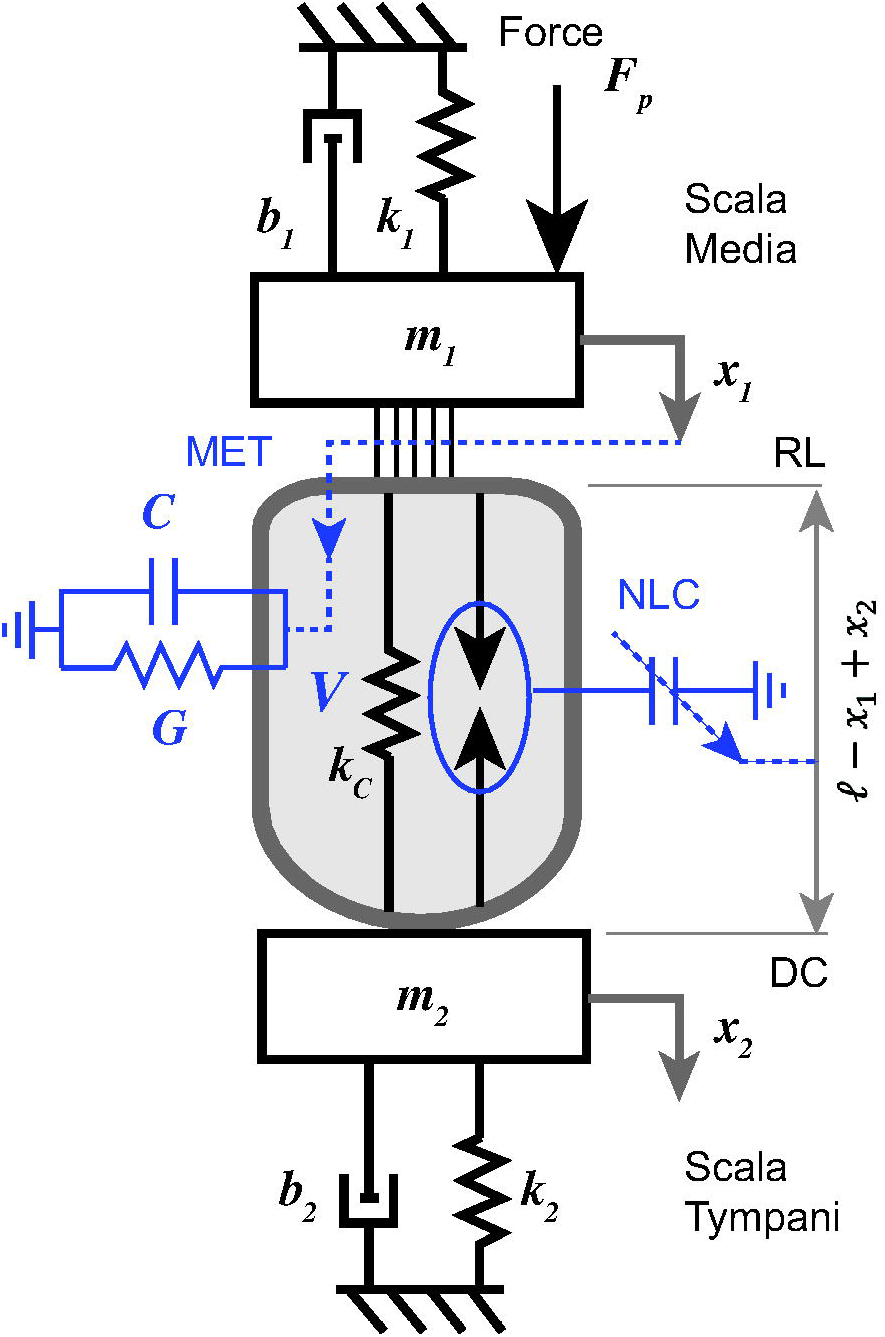
Schematic of the model with a single-compartment piezoelectric OHC loaded by two spring-mass-damper systems.

### Time Domain Equations

The OHC is treated as a space clamped piezoelectric cell undergoing changes in length [42]. The membrane potential *v_m_* is the addition of the resting voltage *v_o_* plus a voltage perturbation *v*. For axisymmetric, isochoric deformations under turgor pressure, the piezoelectric shell equations [47] can be reduced to one dimension relating the OHC axial strain s to the axial force *f_o_* and the transmembrane voltage perturbation *v* according to:

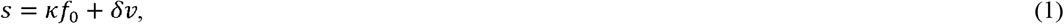

where *κ* (N^−1^) is the axial strain compliance and *δ* (V^−1^) is the axial piezoelectric strain coefficient. The whole-cell axial displacement from the resting length ℓ (m) is approximated as linear using the strain times the length (*x*_1_ – *x*_2_) = *sℓ* (m). Solving for the force in the prestin-motor complex:

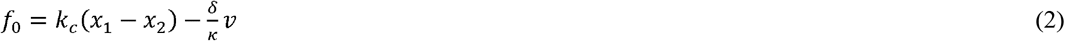

where *x*_1_ and *x*_2_ are displacements of the apical and basal ends of the cell, 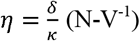 is the piezoelectric force coefficient, and *k_c_* (N-m^−1^) is the axial stiffness of the OHC which is related to the strain compliance *κ* (N^−1^) and the length ℓ (m) by *k_c_* = 1/*κ*ℓ [33]. The mechanical loads acting on the OHC at the apical and basal ends have effective stiffness *k_n_* (N-m^−1^), mass *m_n_* (kg) and viscous drag *b_n_* (N-m^−1^-s^−1^). Newton’s second law for the apical load provides:

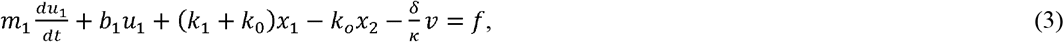

where *f* (N) is the force per OHC arising from the sound-induced dynamic pressure. The velocity at the apical end of the cell is 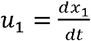 and the velocity at the basal end is 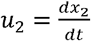. Newton’s second law for the basal load provides:

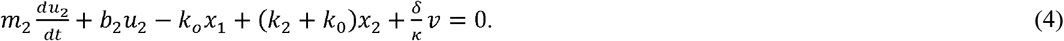

From Kirchhoff’s current law for small deviations in voltage from the resting state (rest: *v_o_, i_o_*), the perturbation in voltage *v* is related to the perturbation in current *i* = *i_M_* + *i_e_* by

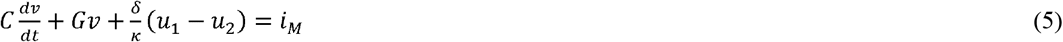

where *C* is the passive electrical membrane capacitance, *G* is the membrane conductance from all ion channels and *i_M_* = *i_MET_* + *i_MB_* is the change in the total mechanically gated current including changes in the MET current from rest and mechano-sensitive basolateral ion channels. NLC arises from piezoelectric coefficient *δ* appearing in the term 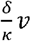 in Eqns. 3-4 and in the term 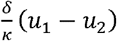 in Eq. 5 (see below). The total mechano-sensitive current is treated as a nonlinear function of apical displacement and velocity, and for simplicity we let the net mechanically gated current be *i_M_* = *i_M_*(*x*_1_, *u*_1_). In this model, the endolymphatic electro-chemical potential is the primary energy source, and cycle-by-cycle modulation of mechanically-gated current provides all of the power for the motor.

The model is minimalistic and is not intended to capture all nonlinearities and details of OHC electro-mechanical power conversion, but is sufficient to reveal some simple relationships between parameters that alter power output. We should note that results in the present report vary the mechanical stiffnesses (*k_n_*). masses (*m_n_*), drag coefficients (*b_n_*), conductance (*G*) and linear capacitance (*C*) in precise ways with cell length (ℓ), but nonlinearities that effect mechanical stiffness and membrane conductance were not included in the simulations. Voltage and strain dependence of the piezoelectric coefficient (NLC, [42]) and nonlinearity of the MET conductance were included to examine the compressive nonlinearity on a single cell level.

### Linearized Matrix Equations

The piezoelectric stress coefficient 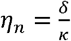 and the mechanically-gated current *i_M_* are nonlinear saturating functions of OHC state and bundle deflection. The total current *i_M_* may include the MET current plus current from strain-sensitive ion channels in the membrane [48]. Mechanical gating of the MET channels in the present model assumes that micromechanics of the organ of Corti directly relates hair bundle deflection to deflection of the apical end of the OHC. For any linear model of the organ of Corti, the displacement of a single hair bundle can be written in the frequency domain as *x_b_* = *X_b_e^jωt^*, and displacement of the apical end of the OHC as *x_b_* = *X_b_e^jωt^*, with the ratio of the two giving the complex-valued, frequency-dependent, transfer function *X_b_*,/*X*_1_. Hence, for small displacements from the resting state, the bundle will deflect with a specific amplitude and phase relative to the apical displacement of the OHC. The linearized model makes no *a priori* assumptions in the present model regarding the how the specific amplitude or phase of the MET current is related to apical hair bundle deflection, and writes the change in the mechanically-gated current in the linearized form: *i_M_* = *σx*_1_ + *αv*_1_. The parameters *σ* and *α* are linearized displacement and velocity gains obtained by optimization (reported in the Results Section). It’s important to note in the frequency domain at CF, the linearized MET current takes the form *I_m_* = (*σ* + *jω_cf_α*)*X*_1_, where the gains (*σ, α*) are cell specific and vary with CF. This means the role of *a* is setting the precise phase of the MET relative to the displacement of the apical cell body. We show in Results that optimum MET phase is close to displacement in the linearized analysis, but the optimum value of *a* is not identically zero. Using the linearized MET current and evaluating the piezoelectric coefficient at the resting voltage reduces Eqns. 2-5 to the matrix form

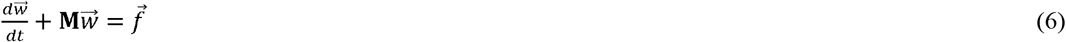

where 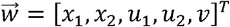, the forcing vector 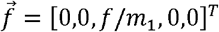. The matrix **M** is

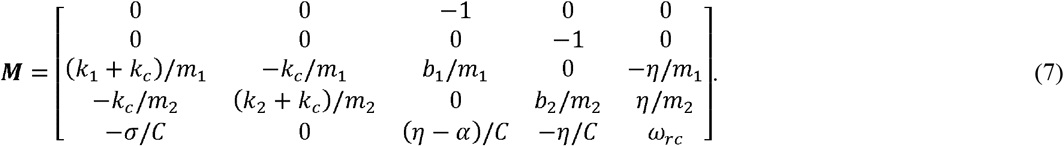

The passive RC corner frequency in the absence of electromotility is defined as *ω_rc_* = *G/C* (rad-s^−1^). The system is stable at the current state if the real part of the eigenvalues of **M** are positive. The solution of Eq. 6 under stable conditions is easily found in the frequency domain using:

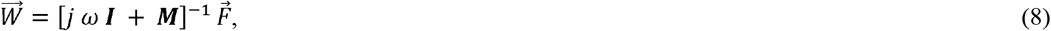

where upper case denotes the frequency domain. The linearized time and frequency domain variables are related by *f*(*t*) = *Fe^jωt^* + *F*e^−jωt^*, *u*(*t*) = *Ue^jωt^* + *U*e^−jωt^* and by *v*(*t*) = *Ve^jωt^* + *V*e^−jωt^*, where the * indicates the complex conjugate.

### Electro-Mechanical Power Conversion

For the nonlinear model, the time-average mechanical power input by the applied force is computed in the time domain using:

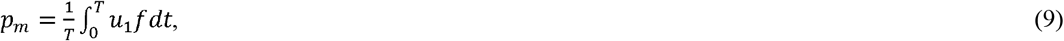

which for small perturbations in the frequency domain becomes 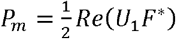. From Eq. 5, the time-average electro-mechanical power conversion by the OHC is:

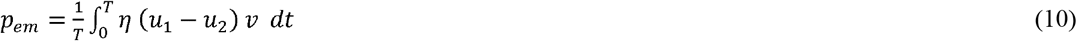

which for small perturbations in the frequency domain becomes 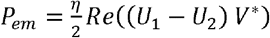. From Eqns. 3-5, the time-average total power delivered by the applied force and the OHC to the viscous load is:

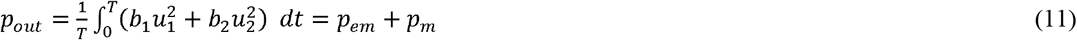

which for small perturbations in the frequency domain becomes 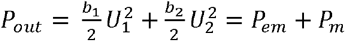. The electrical power lost to OHC basolateral membrane conductance is:

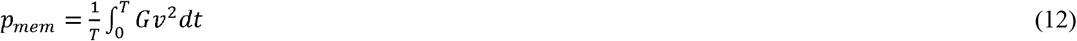

which for small perturbations in the frequency domain becomes 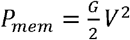.

### Prestin Nonlinearity

The nonlinear capacitance *C_n_* is related to the piezoelectric strain coefficient *δ*, the OHC length ℓ and the axial strain compliance *κ* according to [42]

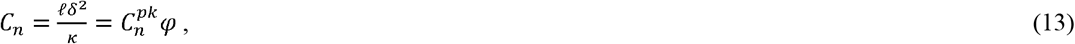

where 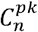 is the peak nonlinear capacitance measured under zero force. The saturating nonlinearity depends on the state of the cell and is defined here for isothermal conditions using a Langevin function

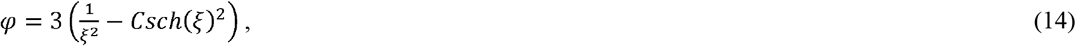

where

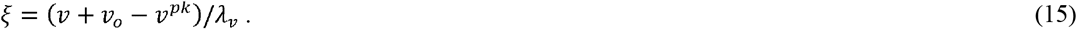

*v_o_* is the resting potential and *v^pk^* is the voltage of peak capacitance under resting stress.

Given the state of the cell at time “*t*”, the nonlinear piezoelectric stress coefficient *η* and piezoelectric strain coefficient *δ* are related to the peak nonlinear capacitance and voltage according to:

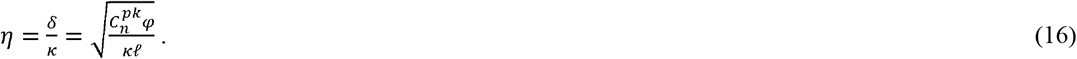

### Nonlinear MET Gating and Current

For time-domain nonlinear simulations, the MET current was modelled using the open probability open probability [49], peak conductance, and electrochemical driving potential. A simple Boltzmann function was used to model the change in the in the open probability: *p*(*z*) = (1 + *e^z/λ^*)^−1^. The change in the MET current relative to the resting current arising from displacement was modeled using:

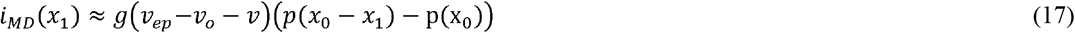

where *g* is the peak conductance, *v_o_* is the resting potential, *v_ep_* is the endolymphatic driving potential, *x_o_* is offset and *λ* is the MET saturation parameter. To relate the parameters in Eq. 17 to the linearized MET displacement gain *σ* described above, we expand Eq. 17 in a Taylor series about *x*_1_ = 0 and set the result to the linearized current *σ x*_1_. The result gives the nonlinear MET current in terms of apical hair cell displacement *x*_1_ and the linearized gain *σ* as:

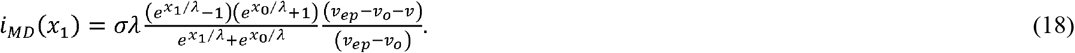

A Taylor series expansion of Eq. 18 confirms that *i_MD_* (*x*_1_) → *σ x*_1_ as *x*_1_ → 0, as required to match the critical point in the linear model. We show in Results that positioning the OHC precisely at the critical point requires a small phase shift in the mechanically gated current that is manifested in the linear model as a velocity gain *α*. To capture this in the nonlinear model we let

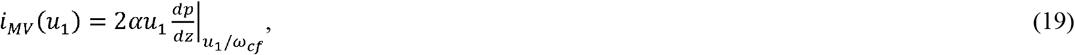

where *i_MV_* → 0 for large velocities (*c.f. u*_1_ > 3 *λω_cf_*). A Taylor series expansion of Eq. 19 confirms that *i_MV_* → *αu*_1_ as *u*_1_ → 0, as required to reproduce the critical point in the linear model. The toatl nonlinear MET current is *i_MD_* + *i_MV_*. Numerical simulations were done with *i_MV_* = 0 and *i_MV_* from Eq. 19 to evaluate the potential importance of MET adaptation in nonlinear power conversion. MET parameters are provided in Table 1. The optimum linearized gains (*σ, α*) required to place the OHC at the critical point for small force stimuli were found analytically (see Results).

**TABLE 1.**
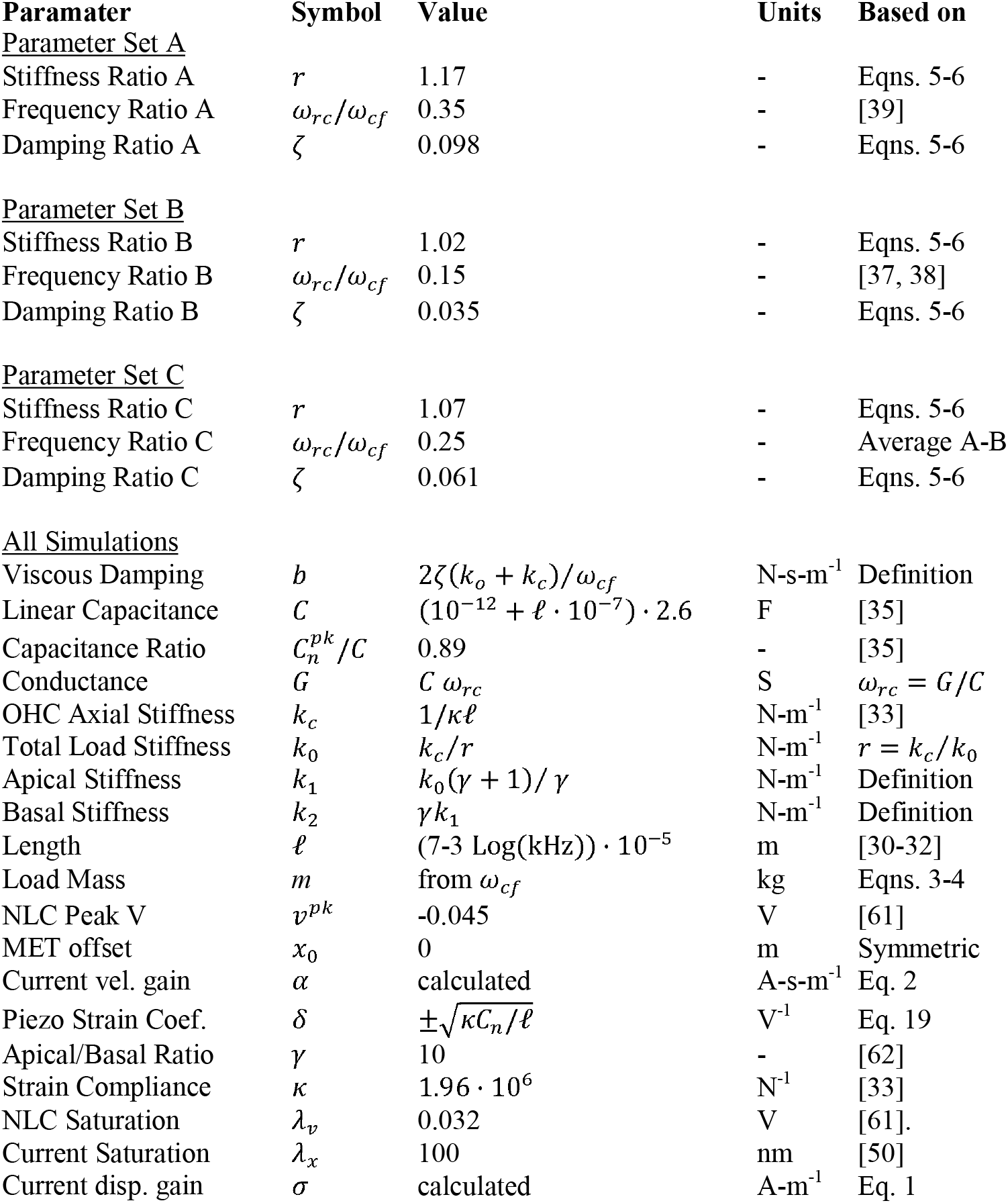
Optimum OHC Parameters.

### Numerical Simulations

Nonlinear numerical simulations (Fig. 2) solved Eqns. 3-5 using a 4^th^ order Runge-Kutta-Fehlberg method with an adjustable step size to maintain a truncation error of <10^−6^ (IgorPro 9, WaveMetrics, Lake Oswego, OR). Nonlinear simulations used the piezoelectric coefficient in Eq. 16 and the MET current in Eq. 18. Linear simulations and stability analysis (Figs. 3–4) were carried out in the frequency domain using Eq. 6. Numerical simulations confirmed that the nonlinear numerical solution matches the frequency domain solution as the applied force approaches zero.

**Fig. 2.**
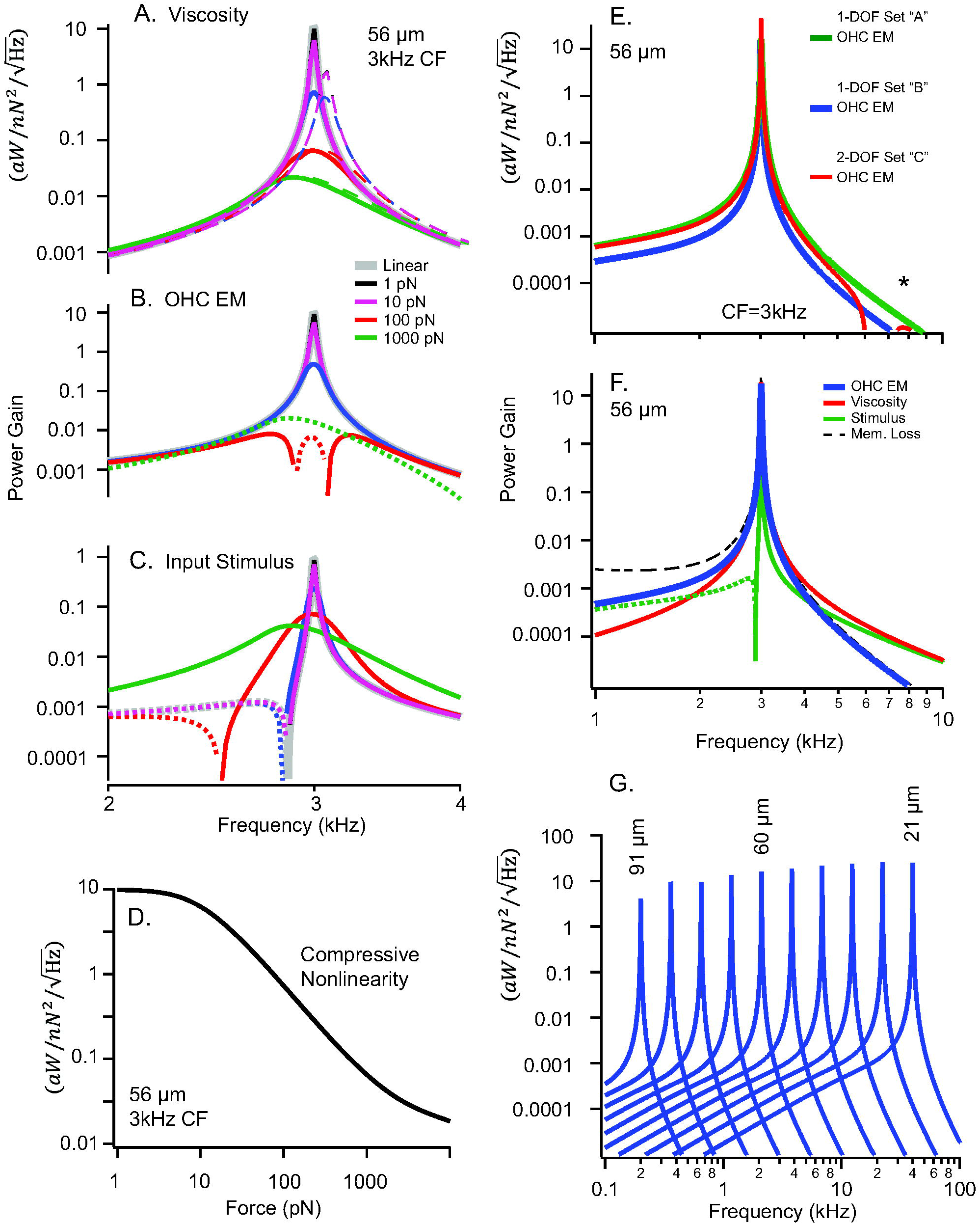
Electro-mechanical power conversion. All curves are normalized by sinusoidal applied force and shown as power gain in units of 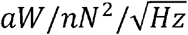. With the exception of panel G, all all results are for a 56 μm long OHC (CF=3kHz). A-C). Predicted power gain for the non-linear model with the magnitude of the force ranging from 1-1000pN. Thick gray curves are from frequency domain linear model, which overlap nonlinear numerical solutions in the time domain for forces of ~5 pN or less. Thick solid curves using an MET current with optimum phase (Eqns. 18-19). A) Power delivered to viscosity (*P_out_*), with thick solid curves using the optimum MET phase and as thin dashed curves aligning the MET phase precisely with apical OHC cell-body displacement (*α* = 0). B) OHC electro-mechanical power conversion (*P_em_*). C) mechanical power associated with the sinusoidal force (*P_m_*). Dotted curves indicate negative power. D) Power gain at CF of 3kHz as a function of sinusoidal stimulus level showing compressive nonlinearity. E-G) Predicted power gain for the linear model. E) Electro-mechanical power conversion *p_em_*) predicted by the 1-DOF linear model using parameter set “A” with *ω_rc_*/*ω_cf_*=0.35 (green, solid) and parameter set “B” with *ω_rc_/ω_cf_*=0.15 (blue, solid). Power conversion predicted by the 2-DOF linear model using parameter set “C” with *ω_rc_/ω_cf_*=0.25 (red, solid). The 2-DOF model shows a second resonance (*), but with a peak ~5 orders of magnitude less than the peak at CF. F) Components of power predicted by the 1-DOF linear model: electro-mechanical power conversion (blue, *P_em_*), power delivered to viscosity (red, *P_out_*), power input by the applied force (green, *P_m_*), electrical power lost to membrane conductance (dashed, *P_mem_*). Dotted curves indicate negative average power. G) Predicted electro-mechanical power conversion under impedance matched conditions for OHCs of various lengths for the 1-DOF model.

**Fig. 3.**
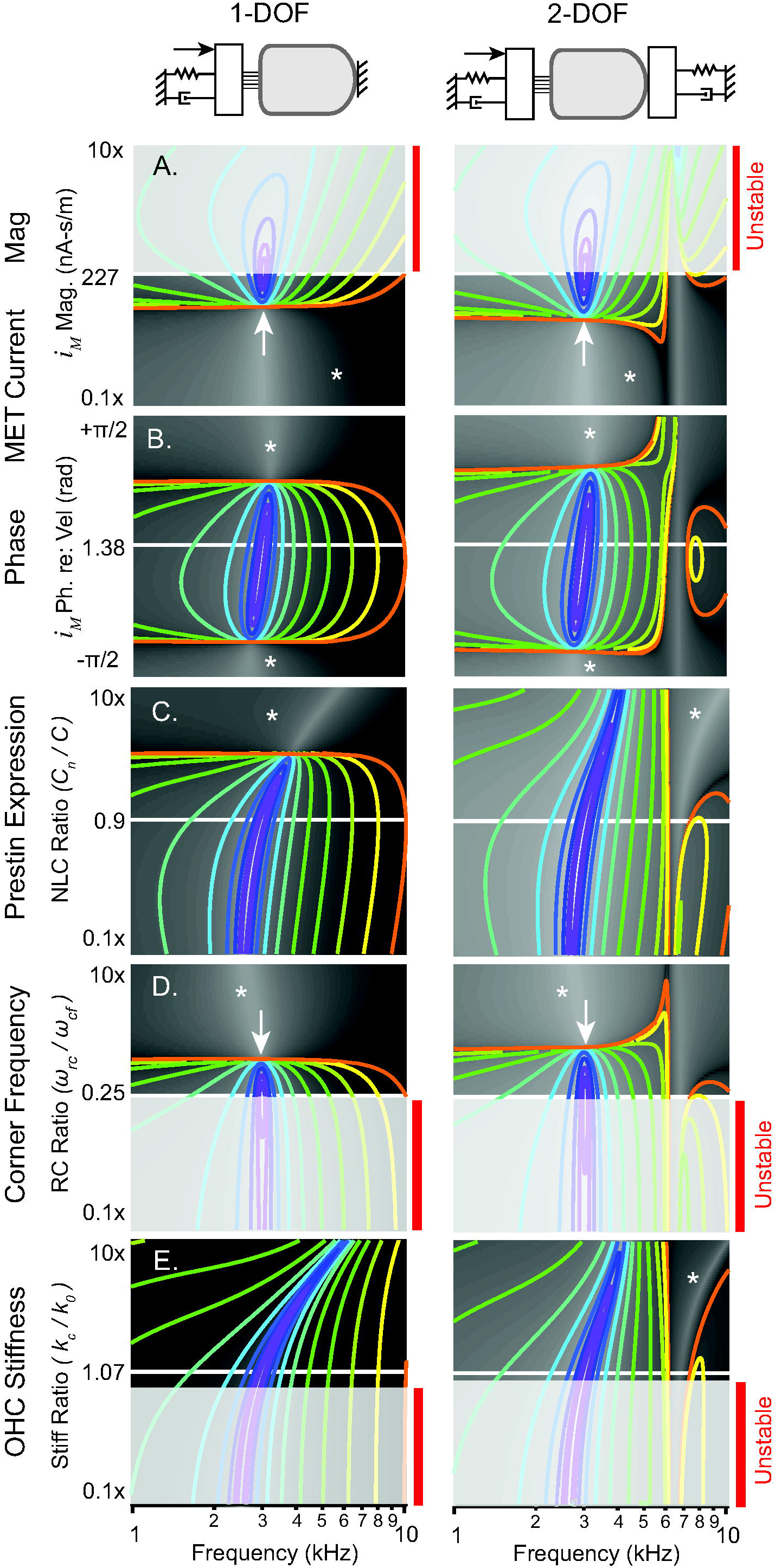
Linear stability and power output. OHC parameters setting the critical point and matching the impedance are the same for 1-DOF (left) and 2-DOF (right) simulations, and give the peak power output for 56 μm long cell located at 3 kHz at the center of each panel. Contour lines show the magnitude of power output, equally spaced on a log scale. OHCs consume mechanical power in regions lacking contour lines (*). Red bars and gray shading indicate regions of limit-cycle oscillation. A-B) Changes in power output and stability when changing MET gain and phase. The critical point is at the center of A and B. C-D) Impedance matching and sensitivity of power output to changes in OHC parameters. C) Sensitivity of power output to capacitance ratio, 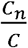. High levels of prestin expression and high *C_n_* stiffen the system and ultimately cause the OHC to consume power rather than output power. D) Sensitivity of power output to the ratio of the RC corner to CF, 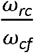. Power output is highly sensitive to *ω_rc_* and stable power output occurs only over a limited range between the white arrow and light gray region. D). Sensitivity of power output to the stiffness of the OHC vs the load, 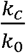. Instability is predicted to occur if the OHC stiffness is too low.

**Fig. 4.**
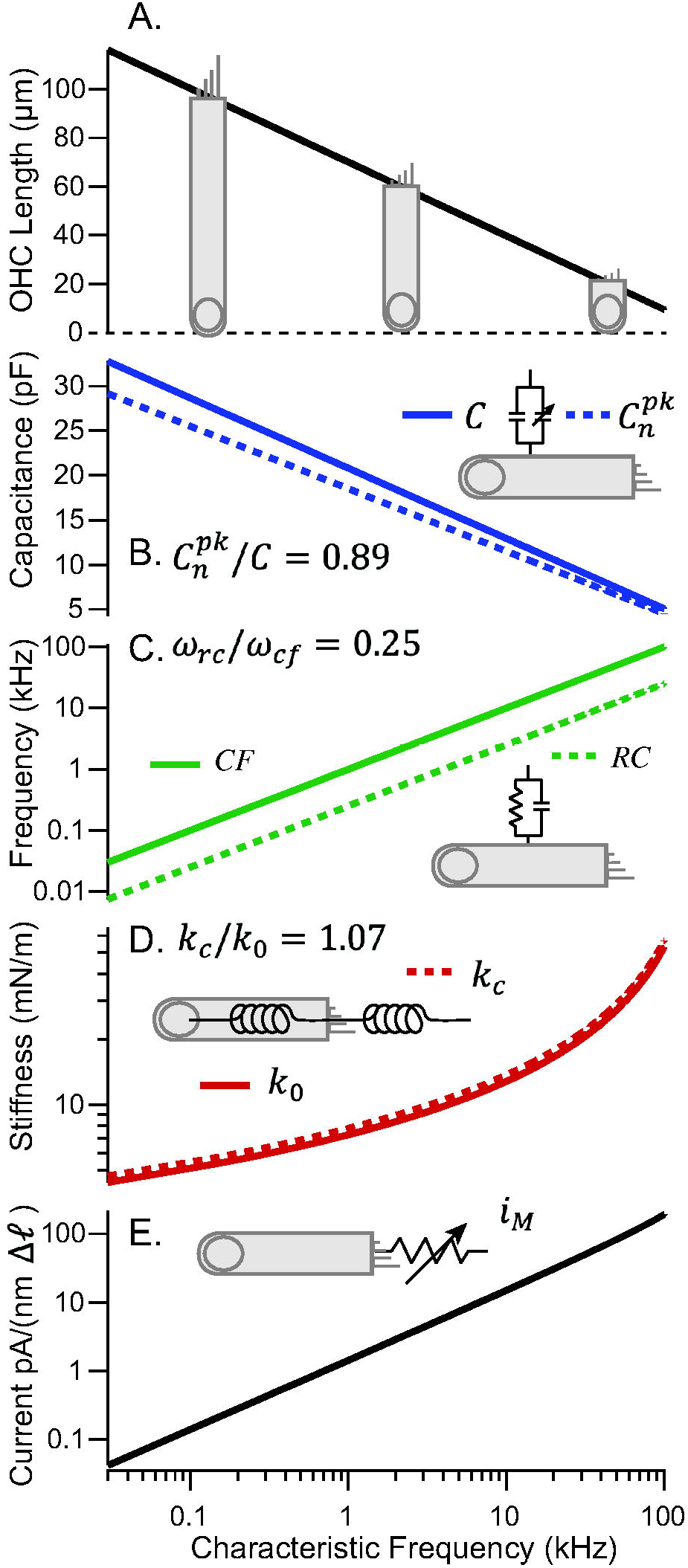
Summary of OHC parameters that are predicted to position the cell at the critical point and optimize electro-mechanical power conversion by impedance matching. A). OHC length vs. CF. B) Optimized linear and nonlinear capacitance. C) Optimized RC corner frequency. D) Optimized OHC stiffness. E) Optimized mechano-sensitive current.

## RESULTS

### Gating of the MET current sets the critical point

The model equations are unstable under certain conditions. Sensitivity of the MET current to displacement is the key factor that determines the bifurcation point where the cell transitions from stable to unstable, and positions the operating point at the edge of dynamic instability to maximize tuning and gain [44–46]. It is sufficient to use the linearized equations to find the MET parameters that place the OHC at the critical point where vibration is maximized. The critical point conditions were found by solving the linearized version of Eq. 6 analytically using symbolic manipulation and finding the values of *σ* and *α* that result in infinite Q resonance at CF (*ω_cf_* rad-s^−1^) (see Supplement Mathematica code). In the linear limit, the OHC operates at the critical point if the displacement gain *σ* is:

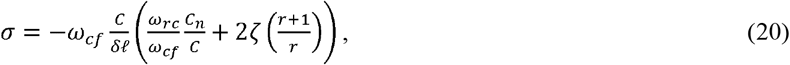

and the velocity gain *α*_0_ is:

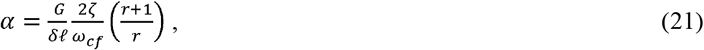

where *C* is the linear capacitance, *C_n_* is the nonlinear capacitance, *δ* is the piezoelectric strain coefficient, ℓ is the cell length, *ω_rc_* = *G/C* is the passive RC corner frequency, *G* is the linearized membrane conductance, 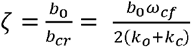 is the nondimensional passive damping ratio and 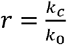 is the stiffness of the cochlear load *k_c_* divided by the OHC axial stiffness *k*_0_.

We call Eq. 20–21 the *critical point conditions* because they maximize tuning and gain by placing the system on edge of instability. This is equivalent to finding the gains that perfectly cancel the dissipative effects of viscosity and membrane conductance. It is significant to note appearance of three nondimensional ratios in these equations: the frequency ratio 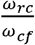, the stiffness ratio 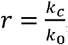, and the capacitance ratio 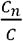. Since the ratios set the critical point, they are *control parameters* in the context of bifurcation theory.

### The critical point conditions are insensitive to the mechanical origins of CF

Two different loading conditions were considered to determine to what extent the critical point conditions depend on mechanics: a 1-DOF load where the apical end of the cell is allowed to move both the basal end of the cell is fixed (*x*_2_ = 0), and a 2-DOF load where both ends of the cell are allowed to move. In both cases, selecting the MET gains using the critical point conditions (Eqns. 20-21) cancels viscous drag and membrane conductance, and sets the characteristic frequency of maximum vibration to the undamped natural frequency of the system. For the 1-DOF model we set *x*_2_ = 0, *x*_1_ = *x*_0_, *k*_1_ = *k*_0_, *m*_1_ = *m*_0_, *b*_1_ = *b*_0_, and solve for Eq. 6 for CF (*ω_cf_*, rad-s^−1^):

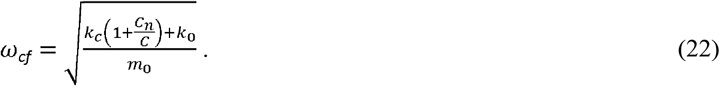

For the 2-DOF model we assume symmetric mass and viscous drag on the two ends of the cell (*m*_1_ = *m*_2_, *b*_1_ = *b*_2_), and write the mechanical stiffness at the basilar membrane in terms of the stiffness at the reticular lamina by *k*_2_ = *γk*_1_. Solving for CF:

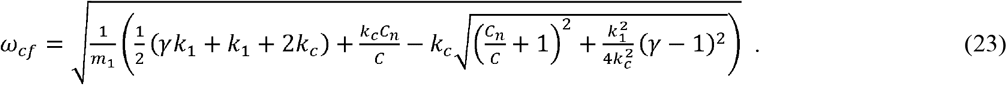

Although equations for CF differ, the critical point conditions (Eqns. 20-21) are exactly the same for the 1-DOF and 2-DOF models providing the compliance of the total load is the same, which requires the stiffness in the 1-DOF model *k*_0_ to be related to the stiffnesses *k*_1_ & *k*_2_ in the 2-DOF model by: 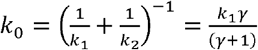. The fact that the critical point conditions are the same for the two models suggests Eqns. 20-21 might be universal, putatively because the nondimensional ratios 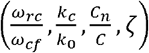 account for the mechanical impedance of a generic load.

### Impedance matching sets optimum OHC parameters

The Q factor of the resonance is theoretically infinite in the linearized equations at CF if the mechano-sensitive gains are set at the critical point (Eqns. 20-21), but not all combinations of parameters result in the same power conversion efficiency. To find optimum parameters with highest efficiency, we minimized the total mechanically-gated current at the critical point with respect to cell length *ℓ* and membrane conductance *G*. In the optimization we approximated the linear and nonlinear capacitance to be proportional to cell length [35], and stiffness to be inversely proportional to cell length [33]. Results of the optimization show the maximum efficiency requires the frequency ratio 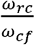 to be related to the stiffness ratio 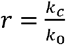 by:

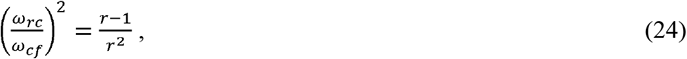

and requires the capacitance ratio 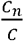 to be related to the damping ratio *ζ* and the stiffness ratio *r* by:

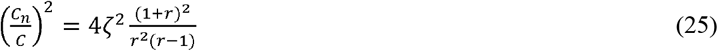

(see Mathematica Supplement for derivation). We call Eqns. 24-25 the *impedance matched conditions*. Since the damping ratio *ζ* is set by passive mechanics, Eqns. 24-25 place strong constraints on the OHC ratios 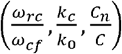, because specifying any one of the three ratios determines the other two. Importantly, like the critical point conditions, the impedance matched conditions are identical for both the 1-DOF and 2-DOF models providing the compliance of the total load is the same in both cases 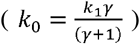. Equivalence suggests the critical point and impedance matched conditions derived here might be general and extend to any generic load.

### Impedance matching conditions are independent of CF

Mammals can manipulate the stiffness of the OHC through genes controlling the cytoarchitecture (right-hand side of Eq. 24), and can manipulate the OHC RC corner frequency though independent genes controlling membrane conductance (left-hand side of Eq. 24). Since the genes effecting the left- and right-hand sides of Eq. 24 are independent variables, we conclude by the mathematical principle of separation of variables (often credited to Wilhjelm Liebniz, 1691) that both the left- and right-hand sides of Eq. 24 are constants, independent of CF and location in the cochlea. This means the ratios 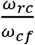 and 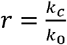 are constants. The same reasoning applies to Eq. 25, and leads to the additional conclusion that the 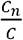 is a constant, independent of CF. It is also significant that the three ratios are the control parameters that set the critical point of maximum tuning and power output (Eqns. 20-21), and therefore are the key parameters that optimize OHCs for cochlear amplification.

### Parameters

Voltage clamp data from guinea pig OHCs reported by Corbitt et al. [35] gives 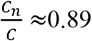, and as predicted by Eq. 24 the ratio is nearly constant (s.d. 0.19; s.e.m. 0.03) for hair cells of various lengths across CF locations (~ 0.1-40 kHz). Two reports from gerbil and guinea pig OHCs suggest the ratio of the RC corner frequency to the CF is near 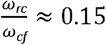 [37, 38], while another report from rat and gerbil OHCs place the ratio near 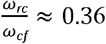 [39] independent of CF. Present simulations therefore consider 3 parameter sets listed in Table 1, corresponding to 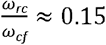, 0.25 and 0.36 (although results show little difference in power output). Taking the ratios 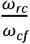 and 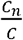 as known constants independent of CF, Eq. 24–25 were solved for the damping ratio 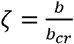 and the stiffness ratio 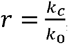, which are also constants independent of CF. For the 2-DOF model 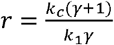. The mass *m* was found from *ω_cf_* (Eq. 22–23). The OHC linear capacitance *C* was found from the cell membrane surface area (Table 1), which also specifies the peak nonlinear capacitance under zero load using the ratio 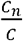. The conductance *G* is found from the capacitance and RC corner frequency. The nonlinear capacitance and compliance provide the piezoelectric strain coefficient *δ* (Eq. 16). In the present simulations we set the mechano-sensitive current gains to 96% of the optimum values specified by Eqns. 19-20, to ensure the system was on the stable side of the critical point. Hence, the constants 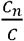 and 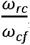, and the CF at the cell location are sufficient to specify all OHC parameters (using Eqns. 20-25).

### Nonlinear Numerical Results

To examine power amplification and the role of nonlinearity, we computed OHC electro-mechanical power conversion as a function of stimulus force amplitude. Parameters were selected as described above and Eqns. 3-5 were solved numerically in the time domain using sinusoidal applied forces including nonlinearities in prestin and the MET (Eqns 13-19). After steady state oscillation was reached, Eqns. 9-12 were applied in the time-domain to compute the time-averaged mechanical power output and electrical power consumed. For low force levels (~<5 pN), numerical solutions of the full nonlinear system are identical to the closed form analytical solution of the linearized model (Eq. 6). Fig. 2A-D show results for the 1-DOF model for a 56μm long cell tuned to 3kHz CF, with the critical point, CF, and impedance match set according to Eqns. 20-25 (Table 1, parameter set “C” used unless otherwise noted). Thick solid curves use the nonlinear piezoelectric coefficient and nonlinear MET current described by Eqns. 13-19. Fig. 2A shows the time-average power delivered to the viscous load for sinusoidal forcing ranging from 1-1000 pN, with results normalized by force and frequency and plotted as power gain 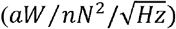. As the applied force becomes small (e.g.2A, 1pN, black curve), the power gain delivered to viscosity in the nonlinear model becomes sharply tuned and equal to the power gain in the linear model (2A, gray curve). The peak power output occurs for a stimulus frequency precisely at the 3kHz CF, and is attenuated ~4 orders of magnitude when the stimulus frequency is reduced to 2kHz. The best frequency decreases as the response transitions from active amplification (2A, black, 1pN) for low force levels to passive vibration for high force levels (2A, green, 1000pN). Dashed curves in Fig. 2A illustrate how the power gain is reduced if the MET phase is fixed to precisely align with OHC displacement (by setting *α* = 0 in Eq. 19) instead of set at the optimum (Eq. 21). The phase shift introduced by *α* in this model cell is only 0.17 radians, yet is predicted to shift the CF up and reduce the peak power gain. The mechanical power delivered to viscosity (2A) for low stimulus force levels (<10 pN) comes primarily from OHC electro-mechanical power conversion (2B, black & purple curves), and to a lesser extent from the applied mechanical force (2C, black & purple curves). As the magnitude of the sinusoidal force is increased, OHC electromechanics begins to consume mechanical power rather than providing power (2B, dotted curves) due to a shift in the phase of the voltage modulation relative to velocity. At frequencies below CF, there is an interplay between OHC electro-mechanics and the applied force, resulting in highly inefficient delivery of power to viscosity (2C, dotted curves). Fig. 2D shows the effect of the OHC nonlinearity on power output at 3kHz CF in the form of power gain vs. applied mechanical force magnitude, illustrating >3 orders of magnitude power amplification at low force levels relative to passive power delivery at high levels.

### Linear Results

We used the linearized model to examine parameter sensitivity, power distribution, and stability. Electro-mechanical power conversion (*P_em_*) is shown in Fig. 2E using the 1-DOF model for parameter set A (solid green, 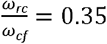) and for parameter set B (solid blue, 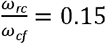). Also shown for comparison is electro-mechanical conversion using the 2-DOF model for parameter set C (solid red, 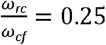). OHC impedance matched conditions (Eqns. 24-25) and critical point conditions (Eqns. 20-21) were identical for 1-DOF and 2-DOF simulations. Similarity between the 3 curves in Fig. 2E is striking, and occurs because the OHC is impedance matched to the total load at 3kHz in all 3 simulations.

To illustrate how the power is distributed, Fig. 2F shows OHC electro-mechanical conversion (*P_em_*, blue), power delivered to viscosity (*P_out_*, red), power supplied by the applied force (*P_m_*, green), and power lost to the membrane conductance (*P_mem_*, black dashed). The dotted green curve indicates the power delevered by the stimulus is negative, but shown on the figure as the absolute value. As required from conservation of energy, the power delivered to viscosity is the addition of the electrical and mechanical terms: *P_out_* = *P_em_* + *P_m_*. To illustrate the role of OHC length and CF, Fig. 2G shows electro-mechanical power conversion by different OHCs along the tonotopic map ranging from 91 to 21 μm in length.

### Parameter Sensitivity and Stability

The linearized model was used to examine system dynamic stability and sensitivity of mechanical power output to changes in MET gains (*σ, α*), capacitance ratio 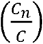. frequency ratio 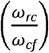 and stiffness ratio 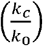. Results in Fig. 3 show changes in stability and power output for a 56μm long OHC tuned to 3kHz CF. In all panels, the horizontal axis is the forcing frequency. Contour lines indicate positive OHC power output with contours equally spaced on a log scale. Regions lacking contour lines (Fig. 3, *) correspond to parameter space where the OHC consumes power rather than outputting power, and regions shaded light grey correspond to parameter space where the system is unstable and the linear model is invalid. Results are shown for the 1-DOF model (left) and the 2-DOF (right), with both models using the same overall compliance of the load 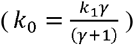, which places the peak power output at the center of each panel for both the 1-DOF and 2-DOF models. The optimum parameters are the same for 1-DOF and 2-DOF models because the critical point conditions (Eq. 20–21) and the impedance matched conditions (Eq. 24–25) are identical.

To position the OHC at the critical point, the mechano-sensitive current must have a specific magnitude and phase relative to the apical cell displacement (Eq. 20–21). Fig. 3A-B illustrate how power output and stability change if the MET current gain (3A) or phase (3B) deviate from the critical point. If the MET gain is too small, the OHC consumes power instead of outputting power (3A,*). As the gain is increased the cell begins to output power (3A, white arrow), reaching a peak at the critical point (3A, center). If the gain is above the critical point the system becomes unstable (3A, light grey & red vertical bars). Numerical solution of the nonlinear equations predicts a self-excited limit-cycle oscillation when operating with parameters in the unstable regions. Stable power output is predicted only for a narrow range of gains, between the white arrow and unstable region (4A). The 1-DOF vs. 2-DOF models have identical critical point gains (Eq. 19–20), but the 2-DOF model shows a modestly expanded region of stability and a sharper roll off for frequencies above CF. The optimum MET phase is shifted ~0.19 rad. relative to cell displacement (3B), and a change in phase reduces power output but does not lead to instability (3B).

Fig. 3. C-D illustrate how altering ratios 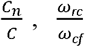, and 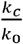 from the impedance matched conditions changes power output and stability. Increasing prestin expression (*C_n_*, Fig. 3C) reduces power output by pushing the best frequency above CF, and if significantly overexpressed can cause the cell to consume power rather than output power (3C, * regions without contour lines). Under-expression of prestin reduces power output primarily by pushing the best frequency below CF. Increasing the RC corner by increasing membrane conductance *G* is predicted to cause the cell to consume power rather than output power (3D, *), while decreasing the RC corner is predicted to induce instability (3D, red vertical bars) and lead to limit cycle oscillation in the nonlinear model. The present analysis predicts stable power output for a relatively narrow range of 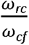, between the white arrow and unstable region (3D). Lowering the OHC axial stiffness below the stiffness of the cochlear partition (per OHC) is also predicted to lead to instability (3E, red vertical bars), while increasing stiffness reduces power output primarily by pushing the best frequency above CF. Power output in the 2-DOF model is relatively insensitive to the relative stiffness at the top of the cell vs. the bottom, providing the combined stiffness is held at the impedance matched value. The primary effect of the stiffness ratio in the 2-DOF model is placement of the anti-resonance frequency and sharpness of the power output roll-off for stimuli above CF.

## DISCUSSION

Our results support the hypothesis that nature has optimized OHC electro-mechanical power conversion at CF by following a universal *parametric blueprint* of OHC design. The blueprint requires setting the MET gains at CF to satisfy the *critical point conditions* (Eqns. 20-21) and setting OHC biophysical parameters based on the nondimensional ratios 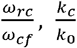 *and* 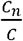 to satisfy the *impedance matching conditions* (Eq. 24–25). Importantly, all three ratios are predicted to be constants, independent of CF. Both the critical point and impedance matching conditions and were found in the present analysis to be exactly the same for two different mechanical loads (1-DOF vs. 2-DOF), suggesting the same conditions might also apply in the cochlea where the load is much more complex. Extending the matching conditions to the cochlea makes specific predictions how OHC properties vary along the tonotopic map, summarized in Fig. 4 (Parameter Set C, Table 1). Given the OHC lengths in Fig. 4A [30–32], the present model predicts universal relationships between the nonlinear capacitance (4B), the passive RC corner frequency (4C), the stiffness (4D), the mechanically gated current (4E-F). As discussed below, all of these predictions compare favorably to available experimental data, supporting the hypothesis that the critical point and impedance matching conditions might apply universally to all OHCs, including in the cochlea.

The prediction that 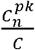 is a universal constant independent of CF is consistent with data in guinea pig [35], where 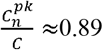 for hair cells of various lengths located from the 0.1-40 kHz regions of the cochlea. Using the linear capacitance based on the cell surface area gives the nonlinear capacitance in Fig. 4B (dotted). The present analysis also predicts that the ratio 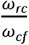 is a universal constant independent of CF, and is likely to fall between 0.15 [37, 38] and 0.35 [39]. Using the average 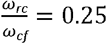 gives the optimum RC corner frequency in Fig. 4D (dotted), which is well below CF (solid). The analysis further predicts that 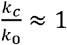 is a universal constant independent of CF. Using the axial strain compliance of OHCs [33] gives the OHC stiffness in Fig. 4D (solid), and since the ratio is a constant also gives the stiffness against which one OHC works in the cochlea (dotted). Stiffness measurements from the in-tact cochlear partition [34] and from isolated OHCs [33] are consistent with this prediction, and suggest both stiffnesses increase with CF roughly in proportion to the inverse of OHC length. The analysis also predicts that the total mechanically-gated current increases with CF (Fig. 4E). The current gain in Fig. 4E is the sum of the MET current plus any mechanically-gated membrane current in the basolateral membrane, and is shown as pA per nm change in OHC length. There are no data currently available to directly test the prediction in Fig. 4E, but there is evidence that the single channel MET current increases substantially with CF [51, 52], and that the expression of the ultrafast mechano-sensitive ion channel K_v_7.4 also increases with CF [48]. We hypothesize the currents combine to provide the net mechano-sensitive current predicted in Fig. 4E.

Present results further suggest the speed of OHC power output is not limited by the speed of the motor itself or the RC corner frequency, but instead is limited by the minimum possible cell length and maximum possible density of ion-channel expression in the basolateral membrane. As CF increases, the OHC length must decrease (Fig. 4A, required to set *k_c_/k*_0_), the basolateral membrane conductance must increase (Fig. 4C, required to set *ω_rc_/ω_cf_*), and the mechanically gated current must increase (Fig. 4E, required to set the critical point). The minimum OHC length feasible to accommodate the nucleus and expression of required membrane proteins is about 8 μm, which places the upper limit of OHC peak power output near 115 kHz (Fig. 4A). Above this frequency, an 8μm long OHC can still output power but efficiency will be sharply attenuated (see Fig. 3G). Interestingly, a minimum OHC length near 8μm is consistent with short OHCs in toothed whales that hear above 100kHz [2, 53], and 115 kHz is close to the upper frequency estimated from behaviour for echolocating killer whales [54].

Optimum power output in this theoretical analysis requires *ω_rc_/ω_cf_* to be a constant, which means the membrane conductance must increase with CF. In order to maintain the resting potential near the point of maximum electromotility (*V^pk^*), the standing current entering the OHC must also increase with CF to counteract the increased conductance. Failure to maintain *ω_rc_/ω_cf_* at the optimum and/or failure to increase the standing current accordingly will lead to a dramatic reduction in power output, or can trigger dynamic instability (Fig. 3D). We hypothesize that the *ω_rc_/ω_cf_* requirement necessitates the silent current observed previously in the cochlea [55], and might explain why the silent current increases in magnitude with CF from apex to base.

Physical parameters that set the mechano-sensitive current to its optimum value position the OHC on the edge of a dynamic instability, a critical point responsible for the sharp tuning and high gain of OHCs under load [44–46]. Here, we used analytical methods to find closed form expressions locating the critical point in parameter space. There are three distinct *control parameters* that can push the system from optimum amplification into limit-cycle oscillation (Fig. 3, red vertical bars, gray regions): increased sensitivity of mechano-sensitive current (gain *σ*), decreased RC corner frequency 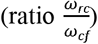, and decreased OHC stiffness 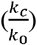. Changes in any of the three control parameters can move the OHC away the critical point, resulting in decreased power output in one direction or limit-cycle instability in the other direction. If a control parameter is changed to cross into an unstable region, the OHC enters a self-sustaining stable limit-cycle oscillation. The limit-cycle frequency of oscillation is below the CF of the cell, about ~5% below for a 56 μm long OHC located at the 3kHz place.

Results demonstrating the RC corner frequency is a control parameter (through action on 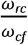) for dynamic stability might have important implications regarding the function of the medial olivocochlear (MOC) efferent system. MOC synaptic contacts on OHCs modulate the conductance of the basolateral membrane (and *ω_rc_*) through nicotinic acetylcholine receptors that trigger opening of Ca^2+^ activated K^+^ channels [56]. Results in Fig. 3D show the OHC is unstable if *ω_rc_* is too low, compelling the hypothesis that tonic firing of MOC efferent neurons sets the OHC at a stable operating point just above the bifurcation, which is the operating point of highest power output. Further increasing MOC efferent activity will push *ω_rc_* above the optimum, eventually into a parameter space where the OHC consumes power rather than outputting power. Function as a control parameter for setting the operating point relative to the critical point gives the MOC system exquisite control over cochlear amplification, with the ability to influence power output well beyond what would be expected from a simple change in receptor potential modulation caused by an efferent mediated change in membrane conductance. This *control parameter* mechanism is likely responsible for the dramatic reduction of mechanical vibrations [57] and reduction in spiral ganglion neuron sensitivity [58, 59] caused by activation of the MOC system. MOC function as a control parameter is also consistent with the influence of contralateral acoustic stimulation on spontaneous otoacoustic emissions (SOAEs). Present results (Fig. 3D) predict a low OHC basolateral conductance can lead to instability and limit cycle oscillation (e.g. Fig. 4F), which would be manifested as a SOAE just below CF. Activation of the MOC by contralateral acoustic stimulation would increase the control parameter 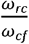 and move the operating point toward stability (Fig. 3D). Our numerical simulations predict MOC activation will reduce the SOAE amplitude and shift the frequency up, exactly as seen experimentally [60].

There are numerous limitations of the present analysis that should be noted. First, the model consists of a very simple isolated OHC driving a linear spring-mass-damper load. No attempt was made to model OHC electro-mechanical power conversion within the complex 3D cochlear environment, and no attempt was made to simulate tuning curves in the cochlea. Using the optimized OHC parameters predicted here (Fig. 4) in a full 3D cochlear model that includes longitudinal coupling and traveling waves would not be expected to generate extremely sharp tuning curves, but would be expected to optimize OHC electro-mechanical power efficiency. Similarly, the sharp tuning for isolated model cells in Figs 2–3 is an artifact of model simplicity and is not possible in reality, but the parameters required to set the critical point and to match the impedance would be expected to hold. Parameter regions of instability in Fig. 3 are also approximate, and may not extend quantitatively under the more complex conditions in the cochlea.

## AUTHOR CONTIRBUTIONS

Conceived the research (RDR, TAB), performed the mathematics (RDR), wrote the code (RDR), drafted and edited the manuscript (RDR, TAB).

## ACKNOWLEDGMENTS

The authors thank W. Brownell, K. Grosh, J.H. Nam and N. Soley and the reviewers for useful comments on portions of the manuscript.

## FUNDING STATEMENT

Funding was provided by the National Institutes on Deafness and other Communication Disorders grant R01 DC006685 (RDR) and the University of Utah (TCB).

## Notes

Conflict of interest: The authors declare no competing financial interests.

### Competing Interest Statement

The authors have declared no competing interest.

### Summary of Updates

Methods appear before Results. MET nonlinearity and Fig. 2 updated.

