## Supplemental Code for "A Parametric Blueprint for Optimum Cochlear Outer Hair Cell Design" (from another version of this preprint: version 1, 2022-10-14)

#### A. 1- DOF Toy Model

Using the piezoelectric constitutive law ($x == (l\,\kappa\, f + l\,\delta\, v)$) for the OHC and Newton's law for the cochlear load ($\left(\left(k - m\,\omega^2\right) + b\, I\, \omega\right) x == -f + fp$) we eliminate force f and find a single equation relating OHC axial displacement "x" to force "fp" applied to the external load and voltage "v" in the OHC. "l" is length, "$\kappa$" is OHC strain compliance (stiffness is ko=1/$\kappa$ l), "m" is total mass of the load, "k" is the added stiffness of the load and "b" is the damping coefficient. "$\omega$" is stimulus frequency and "f" is the force acting on the OHC body.

```
In[∘]:= Eliminate[{x == (l κ f + l δ v), ((k - m ω^2) + b I ω) x == -f + fp}, f]
```

$$\text{Out}[\circ]= \; x - l\,v\,\delta + k\,l\,x\,\kappa + i\,b\,l\,x\,\kappa\,\omega - l\,m\,x\,\kappa\,\omega^2 == fp\,l\,\kappa$$

Solve for the applied force "fp'" in terms of voltage "v" and displacement "x"

$$\left(x - l\,v\,\delta + k\,l\,x\,\kappa + i\,b\,l\,x\,\kappa\,\omega - l\,m\,x\,\kappa\,\omega^2\right) / (l\,\kappa) == fp$$

$$\text{Out}[\circ]= \frac{l\,v\,\delta + b\,l\,u\,\kappa + \frac{i\,u}{\omega} - \frac{i\,k\,l\,u\,\kappa}{\omega} + i\,l\,m\,u\,\kappa\,\omega}{l\,\kappa}$$

Simplify this to the same form given in the manuscript (Eq. A3 for ordinary damping derivative n=1)

$$\left(ko\,x - v\,\frac{\delta}{\kappa} + k\,x + i\,b\,x\,\omega - m\,x\right) = fp$$

$$kp\left(1 - \frac{m\,\omega^2}{(ko + k)} + i\,\frac{b\;\;\omega}{(ko + k)}\right) x - v\,\frac{\delta}{\kappa} = fp$$

where the total stiffness of the partition is the series addition of the load plus the OHC "kp=ko+k", and the non-dimensional damping coefficient "$\zeta$" is

$$\zeta_p = \frac{b\;\;\omega_p}{2\,(k + ko)}$$

We note the effective quality of the passive resonance is $Q = \dfrac{m\,\omega_p}{b} =$

$\frac{m\,\omega_p^2}{\zeta_p\,2\,kp} = \frac{1}{\zeta_p\,2}$. For use in a subsequent section we define the alternative nondimensional damping coefficient with respect to frequency CF

$$\zeta = \frac{b\ \ \omega_{cf}}{2\ (k + ko)} = \zeta_p\,\frac{\omega_{cf}}{\omega_p}$$

$$b = 2\,\zeta_p\,\frac{k + ko}{\omega_p} = 2\,\zeta\,\frac{k + ko}{\omega_{cf}} =$$

The undamped natural frequency of the passive load under voltage clamp conditions is

$$\omega_p = \sqrt{\frac{(k + ko)}{m}}\ .$$

### MET current, OHC velocity and Voltage under Current Clamp and Voltage Clamp

Let the MET current be driven by OHC velocity "u" and OHC displacement "$\frac{u}{I\,\omega}$" according to velocity gain "$\alpha$" and displacement gain "$\sigma$" : Imet = $\left(\alpha + \frac{\sigma}{I\,\omega}\right)$ u. Also include electrode current "a". Solve for velocity "u" and voltage "v"

In[∘]:=
```
Simplify[Solve[
 {((ko + k) - m ω^2 + ⅈ b ω) u/(I ω) - v δ/κ == fp, (I ω c + g) v + δ/κ u == a + (α + σ/(I ω)) u}, {u, v}]]
```

Out[∘]=
$$\left\{\left\{u \to -\frac{\omega\left(\frac{a\,\delta}{\kappa} + fp\,(g + i\,c\,\omega)\right)}{-\frac{\delta\,(i\,\kappa\,\sigma+\delta\,\omega-\alpha\,\kappa\,\omega)}{\kappa^2} + i\,(g + i\,c\,\omega)\,(k + ko + \omega\,(i\,b - m\,\omega))},\right.\right.$$

$$\left.\left. v \to \frac{\kappa\left(fp\,(-i\,\kappa\,\sigma - \delta\,\omega + \alpha\,\kappa\,\omega) - i\,a\,\kappa\,\left(k + ko + i\,b\,\omega - m\,\omega^2\right)\right)}{\delta^2\,\omega + i\,\delta\,\kappa\,(\sigma + i\,\alpha\,\omega) - i\,g\,\kappa^2\,\left(k + ko + i\,b\,\omega - m\,\omega^2\right) + c\,\kappa^2\,\omega\,\left(k + ko + i\,b\,\omega - m\,\omega^2\right)}\right\}\right\}$$

### Resonance under force clamp. Velocity “u” for force clamp (zero current and damping)

In[•]:= $\text{ComplexExpand}\left[-\frac{\omega\,(fp\,(g+i\,c\,\omega))}{-\frac{\delta\,(\delta\,\omega)}{\kappa^2}+i\,(i\,c\,\omega)\,(k+ko+\omega\,(-m\,\omega))}\right]$

Out[•]= $-\frac{fp\,g\,\omega}{-\frac{\delta^2\,\omega}{\kappa^2}-c\,\omega\,\left(k+ko-m\,\omega^2\right)}-\frac{i\,c\,fp\,\omega^2}{-\frac{\delta^2\,\omega}{\kappa^2}-c\,\omega\,\left(k+ko-m\,\omega^2\right)}$

Resonance under force clamp for zero damping occurs when

$-\frac{\delta^2\,\omega}{\kappa^2}-c\,\omega\,\left(k+ko-m\,\omega^2\right)==0$

$-cn-\frac{1}{ko}\,c\,\left(k+ko-m\,\omega^2\right)==0$

In[•]:= $\text{Solve}\left[-cn-\frac{1}{ko}\,c\,\left(k+ko-m\,\omega^2\right)==0,\,\omega\right]$

Out[•]= $\left\{\left\{\omega\to-\frac{\sqrt{c\,k+c\,ko+cn\,ko}}{\sqrt{c}\,\sqrt{m}}\right\},\,\left\{\omega\to\frac{\sqrt{c\,k+c\,ko+cn\,ko}}{\sqrt{c}\,\sqrt{m}}\right\}\right\}$

The resonance frequency in force clamp is

$$\omega_{cf}=\sqrt{\frac{(k+ko)}{m}+\frac{cn}{c}\,\frac{ko}{m}}$$

### Resonance under voltage clamp. The velocity “u” for voltage clamp (zero current and damping)

In[•]:= $\text{Simplify}\left[\text{Solve}\left[\left\{\left((ko+k)-m\,\omega^2\right)\frac{u}{I\,\omega}==\frac{\delta\,v}{\kappa}\right\},\,\{u\}\right]\right]$

Out[•]= $\left\{\left\{u\to\frac{i\,v\,\delta\,\omega}{\kappa\,\left(k+ko-m\,\omega^2\right)}\right\}\right\}$

In[•]:= $\text{Simplify}\left[\text{Solve}\left[k+ko-m\,\omega^2==0,\,\omega\right]\right]$

Out[•]= $\left\{\left\{\omega\to-\frac{\sqrt{k+ko}}{\sqrt{m}}\right\},\,\left\{\omega\to\frac{\sqrt{k+ko}}{\sqrt{m}}\right\}\right\}$

The resonance frequency in voltage clamp is (rad/s)

$$\omega_{vc}=\pm\sqrt{\frac{(k+ko)}{m}}$$

### Resonance under current clamp. The velocity “u” current clamp (zero current and damping)

In[∘]:=
```
ω =.;
fp = 0;
a =.;
```
$$\text{Simplify}\left[\left(-\frac{\omega\left(\frac{a\,\delta}{\kappa}+fp\,(g+i\,c\,\omega)\right)}{-\frac{\delta\,(\delta\,\omega)}{\kappa^2}+i\,(i\,c\,\omega)\,(k+ko+\omega\,(-m\,\omega))}\right)\right]$$

Out[∘]=
$$\frac{a\,\delta\,\kappa}{\delta^2+c\,\kappa^2\left(k+ko-m\,\omega^2\right)}$$

$$\frac{l\,\delta^2}{\kappa}+c\,l\,\kappa\left(k+ko-m\,\omega^2\right)==0$$

$$cn+c\,\frac{1}{ko}\left(k+ko-m\,\omega^2\right)==0$$

In[∘]:=
$$\text{Solve}\left[cn+c\,\frac{1}{ko}\left(k+ko-m\,\omega^2\right)==0,\ \omega\right]$$

Out[∘]=
$$\left\{\left\{\omega\to-\frac{\sqrt{c\,k+c\,ko+cn\,ko}}{\sqrt{c}\,\sqrt{m}}\right\},\left\{\omega\to\frac{\sqrt{c\,k+c\,ko+cn\,ko}}{\sqrt{c}\,\sqrt{m}}\right\}\right\}$$

The resonance frequency in current clamp is

$$\omega_{cc}=\sqrt{\frac{(k+ko)}{m}+\frac{cn}{c}\,\frac{ko}{m}}$$

which is the same as the force clamp undamped resonance frequency

### Find the MET Gains needed to cancel damping at CF

In[∘]:=
```
ω =.;
fp =.;
α =.;
σ =.;
g =.;
a = 0;
Simplify[Solve[
```
$$\left\{\left((ko + k) - m\,\omega^2 + i\,b\,\omega\right)\frac{u}{I\,\omega} - v\,\frac{\delta}{\kappa} == fp,\ (I\,\omega\,c + g)\,v + \frac{\delta}{\kappa}\,u == a + \left(\alpha + \frac{\sigma}{I\,\omega}\right)u\right\},\ \{u, v\}]]$$

Out[∘]=
$$\left\{\left\{u \to -\frac{fp\,\omega\,(g + i\,c\,\omega)}{-\frac{\delta\,(i\,\kappa\,\sigma + \delta\,\omega - \alpha\,\kappa\,\omega)}{\kappa^2} + i\,(g + i\,c\,\omega)\,(k + ko + \omega\,(i\,b - m\,\omega))},\right.\right.$$
$$\left.\left. v \to \frac{fp\,\kappa\,(i\,\kappa\,\sigma + \delta\,\omega - \alpha\,\kappa\,\omega)}{-\delta^2\,\omega + \delta\,\kappa\,(-i\,\sigma + \alpha\,\omega) + i\,g\,\kappa^2\,(k + ko + i\,b\,\omega - m\,\omega^2) + c\,\kappa^2\,\omega\,(-k - ko - i\,b\,\omega + m\,\omega^2)}\right\}\right\}$$

what if we pick MET gains to cancel damping at CF $\omega_{cf} = \sqrt{\frac{(k+ko)}{m} + \frac{cn}{c}\,\frac{ko}{m}}$

The denominator of u is

$$-\frac{\delta\,(i\,\kappa\,\sigma - \alpha\,\kappa\,\omega)}{\kappa^2} + i\,g\,(k + ko + \omega\,(i\,b - m\,\omega)) -$$
$$c\,\omega\,\omega\,i\,b - (c\,\omega\,(k + ko + \omega\,(-m\,\omega)) + cn\,\omega)$$

The last term is zero at CF, leaving the denominator

$$-\frac{\delta\,(i\,\kappa\,\sigma - \alpha\,\kappa\,\omega)}{\kappa^2} + i\,g\,(k + ko + \omega\,(i\,b - m\,\omega)) - c\,\omega\,\omega\,i\,b$$

In[∘]:=
$$\text{ComplexExpand}\left[-\frac{\delta\,(i\,\kappa\,\sigma - \alpha\,\kappa\,\omega)}{\kappa^2} + i\,g\,(k + ko + \omega\,(i\,b - m\,\omega)) - c\,\omega\,\omega\,i\,b\right]$$

Out[∘]=
$$-b\,g\,\omega + \frac{\alpha\,\delta\,\omega}{\kappa} + i\left(g\,k + g\,ko - \frac{\delta\,\sigma}{\kappa} - b\,c\,\omega^2 - g\,m\,\omega^2\right)$$

#### MET gains to cancel damping and force the denominator to zero at CF are

In[∘]:=
$$\text{Solve}\left[\left\{\left(-b\,g\,\omega + \frac{\alpha\,\delta\,\omega}{\kappa}\right) == 0,\ \left(g\,k + g\,ko - \frac{\delta\,\sigma}{\kappa} - b\,c\,\omega^2 - g\,m\,\omega^2\right) == 0\right\},\ \{\sigma, \alpha\}\right]$$

Out[∘]=
$$\left\{\left\{\sigma \to \frac{\kappa\,(g\,k + g\,ko - b\,c\,\omega^2 - g\,m\,\omega^2)}{\delta},\ \alpha \to \frac{b\,g\,\kappa}{\delta}\right\}\right\}$$

Check to make sure the denominator of the velocity is zero with these gains (at CF)

In[∘]:= $\sigma = \frac{\kappa}{\delta}\left(g\left(k+ko-m\,\omega^2\right)-b\,c\,\omega^2\right);$

$\alpha = \frac{\kappa}{\delta}\,b\,g;$

$\omega = \sqrt{\frac{(k+ko)}{m}+\frac{cn}{c}\frac{ko}{m}};$

Simplify$\left[-\frac{\delta\,(i\,\kappa\,\sigma+\delta\,\omega-\alpha\,\kappa\,\omega)}{\kappa^2}+i\,(g+i\,c\,\omega)\,(k+ko+\omega\,(i\,b-m\,\omega))\right]$

Out[∘]= $$\frac{\sqrt{\frac{cn\,ko+c\,(k+ko)}{c\,m}}\left(-\delta^2+cn\,ko\,\kappa^2\right)}{\kappa^2}$$

Yes, the denominator of the velocity is zero, noting $\left(-\delta^2+cn\,ko\,\kappa^2\right)=0$

Double check to make sure the denominator of the voltage also zero with these gains (at CF)

In[∘]:= $\sigma = \frac{\kappa}{\delta}\left(g\left(k+ko-m\,\omega^2\right)-b\,c\,\omega^2\right);$

$\alpha = \frac{\kappa}{\delta}\,b\,g;$

$\omega = \sqrt{\frac{(k+ko)}{m}+\frac{cn}{c}\frac{ko}{m}};$

Simplify$\left[-\delta^2\,\omega+\delta\,\kappa\,(-i\,\sigma+\alpha\,\omega)+i\,g\,\kappa^2\left(k+ko+i\,b\,\omega-m\,\omega^2\right)+c\,\kappa^2\,\omega\left(-k-ko-i\,b\,\omega+m\,\omega^2\right)\right]$

Simplify$[\kappa\,(i\,\kappa\,\sigma+\delta\,\omega-\alpha\,\kappa\,\omega)]$

Out[∘]= $$\sqrt{\frac{cn\,ko+c\,(k+ko)}{c\,m}}\left(-\delta^2+cn\,ko\,\kappa^2\right)$$

Out[∘]= $$\frac{-i\,b\,c\,cn\,ko\,\kappa^3-i\,b\,c^2\,(k+ko)\,\kappa^3-i\,cn\,g\,ko\,m\,\kappa^3+c\,\sqrt{\frac{cn\,ko+c\,(k+ko)}{c\,m}}\,m\,\kappa\left(\delta^2-b\,g\,\kappa^2\right)}{c\,m\,\delta}$$

Yes, the denominator of the voltage is also zero, noting $\left(-\delta^2+cn\,ko\,\kappa^2\right)=0$

Note, the optimum MET gains reduce to:

$$\sigma = \frac{\kappa\left(g\left(k+ko-m\,\omega^2\right)-b\,c\,\omega^2\right)}{\delta}$$

$$\sigma = \frac{\kappa\left(g\left(k+ko-(k+ko)-\frac{cn}{c}\,ko\right)-b\,c\,\omega^2\right)}{\delta}$$

$$\sigma = \frac{\kappa}{\delta}\left(-g\left(\frac{cn}{c}\,ko\right)-b\,c\,\omega^2\right)$$

The corresponding velocity MET gain is

$$\alpha = \frac{b\,g\,\kappa}{\delta}$$

#### Optimum phase of MET

The MET current is: $i=(\alpha\,j\,\omega+\sigma)$*displacement, which gives the Tan(phase) of the current relative to displacement as

In[∘]:= $\text{Simplify}\left[\left(\frac{b\,g\,\kappa}{\delta}\,\omega\right) \Big/ \left(\frac{\kappa}{\delta}\left(-g\left(\frac{cn}{c}\,ko\right) - b\,c\,\omega^2\right)\right)\right]$

Out[∘]= $-\frac{b\,c\,g\,\omega}{cn\,g\,ko + b\,c^2\,\omega^2}$

Define $\zeta$ so $b = 2\,\zeta\,\frac{k+ko}{\omega_{cf}}$ so

In[∘]:= $\text{Simplify}\left[-\frac{2\,\zeta\,(k+ko)\,c\,g}{cn\,g\,ko + 2\,\zeta\,(k+ko)\,c^2\,\omega}\right]$

Out[∘]= $-\frac{2\,c\,g\,(k+ko)\,\zeta}{cn\,g\,ko + 2\,c^2\,(k+ko)\,\zeta\,\omega}$

$$\text{Cot}[\vartheta] = -\frac{\frac{1}{2\,\zeta}\,\frac{cn}{c}\,ko + (k+ko)\,\frac{\omega_{cf}}{\omega_{rc}}}{(k+ko)}$$

$$\text{Cot}[\vartheta] = -\frac{\omega_{cf}}{\omega_{rc}} - \frac{1}{2\,\zeta}\,\frac{cn}{c}\,\frac{ko}{(k+ko)}$$

$$\text{Cot}[\vartheta] = -\frac{\omega_{cf}}{\omega_{rc}} - \frac{1}{2\,\zeta}\,\frac{cn}{c}\,\frac{r}{(1+r)}$$

where $r = \frac{ko}{k}$

### Minimize total MET current, Using MET gains that cancel damping

We are free to pick the relative parameters in the gains. Suppose OHCs are optimized to cancel damping and minimize current for a given velocity . For a unit velocity, the total MET current at CF is $(\alpha + \sigma / j\,\omega)$

$\alpha = \frac{b\,g\,\kappa}{\delta}$

$\sigma = = \frac{\kappa}{\delta}\left(-b\,c\,\omega^2 - g\,ko\,\frac{cn}{c}\right)$

$b = 2\,\zeta\,\frac{k+ko}{\omega_{cf}}$

$$\alpha = \frac{g\,\kappa}{\delta}\,\frac{2\,\zeta}{\omega}\,(k+ko);$$
$$\sigma = \frac{-\kappa}{\delta}\left(g\left(\frac{cn}{c}\,ko\right) + (2\,\zeta\,(k+ko)\,)\,c\,\omega\right);$$
met = α + σ / (I ω)

Out[∘]= $$\frac{2\,g\,(k+ko)\,\zeta\,\kappa}{\delta\,\omega} + \frac{i\,\kappa\left(\frac{cn\,g\,ko}{c} + 2\,c\,(k+ko)\,\zeta\,\omega\right)}{\delta\,\omega}$$

at CF, the magnitude can be written

$$\left(\frac{\kappa\,g\,ko}{\delta\,\omega}\right)^2\left(\left(2\,\zeta\,\frac{(k+ko)}{ko}\right)^2 + \left(\frac{cn}{c} + 2\,\zeta\,\frac{(k+ko)}{ko}\,\frac{\omega_{cf}}{\omega_{rc}}\right)^2\right)$$
$$\left(\frac{\kappa\,g\,ko}{\delta\,\omega}\right)^2\left(\left(2\,\zeta\left(1+\frac{1}{r}\right)\right)^2 + \left(\frac{cn}{c} + 2\,\zeta\left(1+\frac{1}{r}\right)\frac{\omega_{cf}}{\omega_{rc}}\right)^2\right)$$

The magnitude of the MET per unit velocity is proportional to

In[∘]:= $$\text{Simplify}\left[(2\,g\,(k+ko)\,\zeta\,)\,{}^\wedge 2 + \left(-\frac{cn\,g\,ko}{c} + 2\,c\,(k+ko)\,\zeta\,\omega\right){}^\wedge 2\right]$$

Out[∘]= $$4\,g^2\,(k+ko)^2\,\zeta^2 + \left(\frac{cn\,g\,ko}{c} - 2\,c\,(k+ko)\,\zeta\,\omega\right)^2$$

Consider minimization of the current magnitude with respect to OHC length and conductance

$$4\,g^2\,(k+ko)^2\,\zeta^2 + \left(\frac{cn\,g\,ko}{c} - 2\,c\,(k+ko)\,\zeta\,\omega\right)^2$$

Let the ratio cr = cn/c be constant independent of length (e.g. Farrell 2006). Let c and ko depend on length according to: c=ξl, ko=1/(κ l), where ξ and κ are constants.

$$4\,g^2\,(k+1/(\kappa\,l))^2\,\zeta^2 + (cr\,g\,ko - 2\,\xi\,l\,(k+1/(\kappa\,l))\,\zeta\,\omega)^2$$

Optimize total current by finding the minimum with respect to length and conductance

In[∘]:= $$D\left[4\,g^2\,(k+1/(\kappa\,l))^2\,\zeta^2 + (cr\,g\,ko - 2\,\xi\,l\,(k+1/(\kappa\,l))\,\zeta\,\omega)^2,\,l\right]$$
$$D\left[4\,g^2\,(k+1/(\kappa\,l))^2\,\zeta^2 + (cr\,g\,ko - 2\,\xi\,l\,(k+1/(\kappa\,l))\,\zeta\,\omega)^2,\,g\right]$$

Out[∘]= $$-\frac{8\,g^2\,\zeta^2\left(k+\frac{1}{l\,\kappa}\right)}{l^2\,\kappa} + 2\left(cr\,g\,ko - 2\,l\,\zeta\left(k+\frac{1}{l\,\kappa}\right)\xi\,\omega\right)\left(-2\,\zeta\left(k+\frac{1}{l\,\kappa}\right)\xi\,\omega + \frac{2\,\zeta\,\xi\,\omega}{l\,\kappa}\right)$$

Out[∘]= $$8\,g\,\zeta^2\left(k+\frac{1}{l\,\kappa}\right)^2 + 2\,cr\,ko\left(cr\,g\,ko - 2\,l\,\zeta\left(k+\frac{1}{l\,\kappa}\right)\xi\,\omega\right)$$

Minimization with respect to length “l”
gives

$$-\frac{8 g^2 \zeta^2 (k+ko)}{l} ko + 2 (cr\, g\, ko - 2 l \zeta (k+ko) \xi \omega) \left(-2 \zeta (k+ko) \xi \omega + \frac{2 \zeta \xi \omega}{l \kappa}\right) == 0$$
$$-8 g^2 \zeta^2 (k+ko) ko + 2 (cr\, g\, ko - 2 l \zeta (k+ko) \xi \omega) \left(-2 \zeta (k+ko) l \xi \omega + \frac{2 \zeta l \xi \omega}{l \kappa}\right) == 0$$
$$-8 g^2 \zeta^2 (k+ko) ko + 2 (cr\, g\, ko - 2 \zeta (k+ko) c \omega) (-2 \zeta (k+ko) c \omega + 2 \zeta c \omega ko) == 0$$
$$-8 \zeta^2 (k+ko) ko + 2 \left(cr\, ko - 2 \zeta (k+ko) \frac{c}{g} \omega\right) \left(-2 \zeta (k+ko) \frac{c}{g} \omega + 2 \zeta \frac{c}{g} \omega ko\right) == 0$$
$$-8 \zeta^2 (k+ko) ko + 2 (cr\, ko - 2 \zeta (k+ko) wr) (-2 \zeta (k+ko) wr + 2 \zeta wr\, ko) == 0$$

Minimization with respect to conductance “g” gives

$$8 g \zeta^2 \left(k + \frac{1}{l \kappa}\right)^2 + 2 cr\, ko \left(cr\, g\, ko - 2 l \zeta \left(k + \frac{1}{l \kappa}\right) \xi \omega\right) == 0$$
$$8 \zeta^2 (k+ko)^2 + 2 cr\, ko (cr\, ko - 2 \zeta (k+ko) wr) == 0$$

In[∘]:=

```
Simplify[
 Solve[{-8 ζ^2 (k + ko) ko + 2 (cr ko - 2 ζ (k + ko) wr) (-2 ζ (k + ko) wr + 2 ζ wr ko) == 0,
   8 ζ^2 (k + ko)^2 + 2 cr ko (cr ko - 2 ζ (k + ko) wr) == 0}, {wr, cr}]]
```

Out[∘]=

$$\left\{\left\{wr \to -\frac{ko}{\sqrt{k(-k+ko)}}, cr \to -\frac{2 k (k+ko) \zeta}{ko \sqrt{k(-k+ko)}}\right\}, \left\{wr \to \frac{ko}{\sqrt{k(-k+ko)}}, cr \to \frac{2 k (k+ko) \zeta}{ko \sqrt{k(-k+ko)}}\right\}\right\}$$

In[∘]:=

$$\text{Simplify}\left[\frac{\left(\frac{k^2}{\sqrt{-k (k-ko)}} + \frac{k\, ko}{\sqrt{-k (k-ko)}}\right)}{ko}\right]$$

Out[∘]=

$$\frac{k (k+ko)}{ko \sqrt{k(-k+ko)}}$$

$$\left(\frac{\omega_{cf}}{\omega_{rc}}\right)^2 = \frac{ko^2}{-k^2 + k\, ko}$$

$$\left(\frac{\omega_{cf}}{\omega_{rc}}\right)^2 = \frac{r^2}{r-1}$$

$$\frac{c_n}{c} = 2 \zeta \frac{k (k+ko)}{ko \sqrt{k(-k+ko)}}$$

$$\frac{c_n}{c} = 2 \zeta \frac{(1+r)}{r \sqrt{(r-1)}}$$

$$\left(\frac{c_n}{c}\right)^2 = 4 \zeta^2 \frac{(1+r)^2}{r^2 (r-1)}$$

### Determine r and z

From Johnson data $\frac{\omega_{rc}}{\omega_{cf}} = 0.35$, From Corbit data $\frac{C_n}{C} = 0.89$

In[∘]:=
```
ξ =.;
wr = 0.35;
cr = 0.89;
NSolve[{wr^2 == (r - 1)/r^2, cr^2 == 4 ξ^2 (1 + r)^2/(r^2 (r - 1))}, {r, ξ}]
```

Out[∘]= {{r → 6.9965, ζ → -0.953432}, {r → 6.9965, ζ → 0.953432}, {r → 1.16676, ζ → -0.0978549}, {r → 1.16676, ζ → 0.0978549}}

From Housley data $\frac{\omega_{rc}}{\omega_{cf}} = 0.15$, From Corbit data $\frac{C_n}{C} = 0.89$

In[∘]:=
```
ξ =.;
wr = 0.15;
cr = 0.89;
NSolve[{wr^2 == (r - 1)/r^2, cr^2 == 4 ξ^2 (1 + r)^2/(r^2 (r - 1))}, {r, ξ}]
```

Out[∘]= {{r → 43.4209, ζ → -2.8331}, {r → 43.4209, ζ → 2.8331}, {r → 1.02357, ζ → -0.0345597}, {r → 1.02357, ζ → 0.0345597}}

Try $\frac{\omega_{rc}}{\omega_{cf}} = 0.25$, From Corbit data $\frac{C_n}{C} = 0.89$

In[∘]:=
```
ξ =.;
wr = 0.25;
cr = 0.89;
NSolve[{wr^2 == (r - 1)/r^2, cr^2 == 4 ξ^2 (1 + r)^2/(r^2 (r - 1))}, {r, ξ}]
```

Out[∘]= {{r → 14.9282, ζ → -1.5565}, {r → 14.9282, ζ → 1.5565}, {r → 1.0718, ζ → -0.0616847}, {r → 1.0718, ζ → 0.0616847}}

current is proportional to $2\,\zeta\left(1 + \frac{1}{r}\right)$ -- so the second root is smaller current

In[∘]:=
```
N[2 * 0.95 * (1 + 1/7)]
N[2 * 0.097 * (1 + 1/1.17)]
```

Out[∘]= 2.17143

Out[∘]= 0.359812

**If passive zeta is about 0.07, then zeta in this model would be about 0.2**

In[∘]:=
```
ξ = 0.1;
wr =.;
cr = 0.89;
```

$$\text{NSolve}\left[\left\{wr^2 == \frac{r-1}{r^2},\ cr^2 == 4\,\xi^2\,\frac{(1+r)^2}{r^2\,(r-1)}\right\},\ \{wr, r\}\right]$$

Out[∘]=
```
{{r → 1.17327, wr → -0.354781}, {r → 1.17327, wr → 0.354781},
 {r → -0.0613836 - 0.198175 i, wr → -4.62739 - 1.91655 i},
 {r → -0.0613836 - 0.198175 i, wr → 4.62739 + 1.91655 i},
 {r → -0.0613836 + 0.198175 i, wr → -4.62739 + 1.91655 i},
 {r → -0.0613836 + 0.198175 i, wr → 4.62739 - 1.91655 i}}
```

### B. 2 DOF Model

Recall definition of nonlinear capacitance under zero load

$$C_n = l\,\frac{\delta^2}{\kappa}$$

$$\frac{\delta^2}{\kappa^2} = \text{ko cn}$$

#### Apical and Basal Stiffnesses of the load

Require the apical and basal stiffness to combine in series during low frequency vibration to give the same transverse stiffness of the organ of Corti as used for the 1 DOF model. The total load stiffness is 1/ktotal=1/k1+1/(k1*γ).

Given the OHC stiffness ko, the basal stiffness k2=γ k, and the ratio of the total stiffness to the OHC stiffness r=, the apical stiffness k1 must be

In[∘]:= **Solve[1 / k1 + 1 / (k1 * γ) == 1 / (r * ko), k1]**

Out[∘]= $\left\{\left\{k1 \to \frac{\text{ko r } (1+\gamma)}{\gamma}\right\}\right\}$

In the following we call k1 the variable k, but use the above formula in numerical results when relating k1 to the OHC stiffness ko, the ratio r and the ratio γ

#### 2 DOF Undamped free vibration

Symmetric forcing case

In[∘]:=
```
ω =.;
b = 0;
g = 0;
σ = 0;
α = 0;
k1 = k;
k2 = k γ;
m1 = m;
m2 = m;
```

$$\text{Simplify}\Big[\text{Solve}\Big[\Big\{\left((ko + k1) - m1\,\omega^2 + i\,b\,\omega\right)\frac{u1}{I\,\omega} - ko\,\frac{u2}{I\,\omega} - v\,\frac{\delta}{\kappa} == fp/2,$$
$$\left((ko + k2) - m2\,\omega^2 + i\,b\,\omega\right)\frac{u2}{I\,\omega} - ko\,\frac{u1}{I\,\omega} + v\,\frac{\delta}{\kappa} == fp/2,$$
$$(I\,\omega\,c + g)\,v + \frac{\delta}{\kappa}(u1 - u2) == \left(\alpha + \frac{\sigma}{I\,\omega}\right)(u1)\Big\}, \{u1, u2, v\}\Big]\Big]$$

Out[∘]=

$$\Big\{\Big\{u1 \to \frac{i\,fp\,\omega\left(2\,\delta^2 + c\,\kappa^2\left(2\,ko + k\,\gamma - m\,\omega^2\right)\right)}{2\left(c\,k^2\,\gamma\,\kappa^2 + k\,(1+\gamma)\left(\delta^2 + c\,\kappa^2\left(ko - m\,\omega^2\right)\right) + m\,\omega^2\left(-2\,\delta^2 + c\,\kappa^2\left(-2\,ko + m\,\omega^2\right)\right)\right)},$$
$$u2 \to \frac{i\,fp\,\omega\left(2\,\delta^2 + c\,\kappa^2\left(k + 2\,ko - m\,\omega^2\right)\right)}{2\left(c\,k^2\,\gamma\,\kappa^2 + k\,(1+\gamma)\left(\delta^2 + c\,\kappa^2\left(ko - m\,\omega^2\right)\right) + m\,\omega^2\left(-2\,\delta^2 + c\,\kappa^2\left(-2\,ko + m\,\omega^2\right)\right)\right)},$$
$$v \to -\frac{fp\,k\,(-1+\gamma)\,\delta\,\kappa}{2\left(c\,k^2\,\gamma\,\kappa^2 + k\,(1+\gamma)\left(\delta^2 + c\,\kappa^2\left(ko - m\,\omega^2\right)\right) + m\,\omega^2\left(-2\,\delta^2 + c\,\kappa^2\left(-2\,ko + m\,\omega^2\right)\right)\right)}\Big\}\Big\}$$

asymmetric forcing case

```
ω =.;
b = 0;
g = 0;
σ = 0;
α = 0;
k1 = k;
k2 = k γ;
m1 = m;
m2 = m;
```

$$\text{Simplify}\Big[\text{Solve}\Big[\Big\{\left((\text{ko} + \text{k1}) - \text{m1}\,\omega^2 + \mathrm{i}\,\text{b}\,\omega\right)\frac{\text{u1}}{\text{I}\,\omega} - \text{ko}\,\frac{\text{u2}}{\text{I}\,\omega} - \text{v}\,\frac{\delta}{\kappa} == \text{fp},$$
$$\left((\text{ko} + \text{k2}) - \text{m2}\,\omega^2 + \mathrm{i}\,\text{b}\,\omega\right)\frac{\text{u2}}{\text{I}\,\omega} - \text{ko}\,\frac{\text{u1}}{\text{I}\,\omega} + \text{v}\,\frac{\delta}{\kappa} == 0,$$
$$(\text{I}\,\omega\,\text{c} + \text{g})\,\text{v} + \frac{\delta}{\kappa}(\text{u1} - \text{u2}) == \left(\alpha + \frac{\sigma}{\text{I}\,\omega}\right)(\text{u1})\Big\}, \{\text{u1}, \text{u2}, \text{v}\}\Big]\Big]$$

Out[∘]=

$$\Big\{\Big\{\text{u1} \to \frac{\mathrm{i}\,\text{fp}\,\omega\left(2\,\delta^2 + \text{c}\,\kappa^2\left(2\,\text{ko} + \text{k}\,\gamma - \text{m}\,\omega^2\right)\right)}{2\left(\text{c}\,\text{k}^2\,\gamma\,\kappa^2 + \text{k}\,(1+\gamma)\left(\delta^2 + \text{c}\,\kappa^2\left(\text{ko} - \text{m}\,\omega^2\right)\right) + \text{m}\,\omega^2\left(-2\,\delta^2 + \text{c}\,\kappa^2\left(-2\,\text{ko} + \text{m}\,\omega^2\right)\right)\right)},$$
$$\text{u2} \to \frac{\mathrm{i}\,\text{fp}\,\omega\left(2\,\delta^2 + \text{c}\,\kappa^2\left(\text{k} + 2\,\text{ko} - \text{m}\,\omega^2\right)\right)}{2\left(\text{c}\,\text{k}^2\,\gamma\,\kappa^2 + \text{k}\,(1+\gamma)\left(\delta^2 + \text{c}\,\kappa^2\left(\text{ko} - \text{m}\,\omega^2\right)\right) + \text{m}\,\omega^2\left(-2\,\delta^2 + \text{c}\,\kappa^2\left(-2\,\text{ko} + \text{m}\,\omega^2\right)\right)\right)},$$
$$\text{v} \to -\frac{\text{fp}\,\text{k}\,(-1+\gamma)\,\delta\,\kappa}{2\left(\text{c}\,\text{k}^2\,\gamma\,\kappa^2 + \text{k}\,(1+\gamma)\left(\delta^2 + \text{c}\,\kappa^2\left(\text{ko} - \text{m}\,\omega^2\right)\right) + \text{m}\,\omega^2\left(-2\,\delta^2 + \text{c}\,\kappa^2\left(-2\,\text{ko} + \text{m}\,\omega^2\right)\right)\right)}\Big\}\Big\}$$

Undamped 2 DOF natural frequencies are when the denominator goes to zero

In[∘]:=

$$\text{Simplify}\Big[\text{Solve}\Big[2\left(\text{c}\,\text{k}^2\,\gamma\,\kappa^2 + \text{k}\,(1+\gamma)\left(\delta^2 + \text{c}\,\kappa^2\left(\text{ko} - \text{m}\,\omega^2\right)\right) + \text{m}\,\omega^2\left(-2\,\delta^2 + \text{c}\,\kappa^2\left(-2\,\text{ko} + \text{m}\,\omega^2\right)\right)\right) == 0, \omega\Big]\Big]$$

Out[∘]=

$$\Big\{\Big\{\omega \to -\frac{\sqrt{\frac{2\,\delta^2 + \text{c}\,\text{k}\,\kappa^2 + 2\,\text{c}\,\text{ko}\,\kappa^2 + \text{c}\,\text{k}\,\gamma\,\kappa^2 - \sqrt{4\,\delta^4 + 8\,\text{c}\,\text{ko}\,\delta^2\,\kappa^2 + \text{c}^2\left(4\,\text{ko}^2 + \text{k}^2\,(-1+\gamma)^2\right)\kappa^4}}{\text{c}\,\text{m}\,\kappa^2}}}{\sqrt{2}}\Big\},$$
$$\Big\{\omega \to \frac{\sqrt{\frac{2\,\delta^2 + \text{c}\,\text{k}\,\kappa^2 + 2\,\text{c}\,\text{ko}\,\kappa^2 + \text{c}\,\text{k}\,\gamma\,\kappa^2 - \sqrt{4\,\delta^4 + 8\,\text{c}\,\text{ko}\,\delta^2\,\kappa^2 + \text{c}^2\left(4\,\text{ko}^2 + \text{k}^2\,(-1+\gamma)^2\right)\kappa^4}}{\text{c}\,\text{m}\,\kappa^2}}}{\sqrt{2}}\Big\},$$
$$\Big\{\omega \to -\frac{\sqrt{\frac{2\,\delta^2 + \text{c}\,\text{k}\,\kappa^2 + 2\,\text{c}\,\text{ko}\,\kappa^2 + \text{c}\,\text{k}\,\gamma\,\kappa^2 + \sqrt{4\,\delta^4 + 8\,\text{c}\,\text{ko}\,\delta^2\,\kappa^2 + \text{c}^2\left(4\,\text{ko}^2 + \text{k}^2\,(-1+\gamma)^2\right)\kappa^4}}{\text{c}\,\text{m}\,\kappa^2}}}{\sqrt{2}}\Big\},$$
$$\Big\{\omega \to \frac{\sqrt{\frac{2\,\delta^2 + \text{c}\,\text{k}\,\kappa^2 + 2\,\text{c}\,\text{ko}\,\kappa^2 + \text{c}\,\text{k}\,\gamma\,\kappa^2 + \sqrt{4\,\delta^4 + 8\,\text{c}\,\text{ko}\,\delta^2\,\kappa^2 + \text{c}^2\left(4\,\text{ko}^2 + \text{k}^2\,(-1+\gamma)^2\right)\kappa^4}}{\text{c}\,\text{m}\,\kappa^2}}}{\sqrt{2}}\Big\}\Big\}$$

$$\sqrt{\frac{2\,cn\,ko + c\,(k + 2\,ko + k\,\gamma) \pm \sqrt{8\,c\,cn\,ko^2 + 4\,cn^2\,ko^2 + c^2\left(4\,ko^2 + k^2\,(-1+\gamma)^2\right)}}{2\,c\,m}}$$

$$\omega_{cf} = \sqrt{\frac{\frac{cn}{c}\,ko + \frac{1}{2}\,(k + 2\,ko + k\,\gamma) \pm ko\,\sqrt{2\,\frac{cn}{c} + \frac{cn^2}{c^2} + \left(1 + \frac{k^2}{ko^2}\,\frac{(\gamma-1)^2}{4}\right)}}{m}}$$

$$\omega_{cf} = \sqrt{\frac{\frac{cn}{c}\,ko + \frac{1}{2}\,(k + 2\,ko + k\,\gamma) \pm ko\,\sqrt{\left(\frac{cn}{c} + 1\right)^2 + \frac{k^2}{ko^2}\,\frac{(\gamma-1)^2}{4}}}{m}}$$

The second frequeny with the "+" sign is the relevant one because it is the one that deforms the OHC the most and has the most power output.

Note, if $\gamma \to \infty$

$$\omega_{cf} \to \sqrt{\frac{\frac{cn}{c}\,ko + \frac{1}{2}\,(k + 2\,ko + k\,\gamma) \pm ko\,\frac{k\,(\gamma-1)}{ko\,2}}{m}}$$

$$\omega_{cf} \to \sqrt{\frac{ko\left(\frac{cn}{c} + 1\right) + k}{m}}$$

which reporduces the 1 DOF result

#### Find MET to perfectly cancel damping in 2 DOF Model

Symmetric forcing case

In[∘]:=
```
ω =.;
b =.;
g =.;
σ =.;
α =.;
k1 = k;
k2 = k γ;
m1 = m;
m2 = m;
Simplify[Solve[{( (ko + k1) - m1 ω^2 + ⅈ b ω) u1/(I ω) - ko u2/(I ω) - v δ/κ == fp / 2,
   ( (ko + k2) - m2 ω^2 + ⅈ b ω) u2/(I ω) - ko u1/(I ω) + v δ/κ == fp / 2,
   (I ω c + g) v + δ/κ (u1 - u2) == (α + σ/(I ω)) (u1)}, {u1, u2, v}]]
```

Out[∘]=

$$\Big\{\Big\{u1 \to -\Big(\Big(i\, fp\, \delta\, \omega \left(2\, \delta^2\, \omega + c\, \kappa^2\, \omega\, (2\, ko + k\, \gamma + \omega\, (i\, b - m\, \omega)) + g\, \kappa^2\, (-2\, i\, ko - i\, k\, \gamma + \omega\, (b + i\, m\, \omega))\right)\Big)\Big/$$
$$\Big(2\, \Big(\delta\, \left(-i\, g\, ko\, \kappa^2 + \left(\delta^2 + c\, ko\, \kappa^2\right) \omega\right)\, (-k + \omega\, (-i\, b + m\, \omega)) - \delta\, (k\, \gamma + \omega\, (i\, b - m\, \omega))$$
$$\left(\delta\, (i\, \kappa\, \sigma + \delta\, \omega - \alpha\, \kappa\, \omega) - i\, \kappa^2\, (g + i\, c\, \omega)\, (k + ko + \omega\, (i\, b - m\, \omega))\right)\Big)\Big)\Big),$$
$$u2 \to \Big(fp\, \omega\, \Big(2\, i\, \delta^2\, \omega - 2\, \delta\, \kappa\, (\sigma + i\, \alpha\, \omega) + g\, \kappa^2\, (k + 2\, ko + \omega\, (i\, b - m\, \omega)) +$$
$$i\, c\, \kappa^2\, \omega\, (k + 2\, ko + \omega\, (i\, b - m\, \omega))\Big)\Big)\Big/$$
$$\Big(2\, \Big(c\, k^2\, \gamma\, \kappa^2\, \omega + \omega\, (-i\, b + m\, \omega)\, \left(-2\, \delta^2\, \omega + \delta\, \kappa\, (-i\, \sigma + \alpha\, \omega) + c\, \kappa^2\, \omega \left(-2\, ko - i\, b\, \omega + m\, \omega^2\right)\right) -$$
$$i\, g\, \kappa^2\, \left(k^2\, \gamma + k\, (1 + \gamma)\, \left(ko + i\, b\, \omega - m\, \omega^2\right) + \omega\, (-i\, b + m\, \omega)\, \left(-2\, ko - i\, b\, \omega + m\, \omega^2\right)\right) +$$
$$k\, \Big(\omega\, \left(\delta^2 + c\, \kappa^2\, \left(ko + i\, b\, \omega - m\, \omega^2\right)\right) +$$
$$\gamma\, \left(\delta^2\, \omega + i\, \delta\, \kappa\, (\sigma + i\, \alpha\, \omega) + c\, \kappa^2\, \omega\, \left(ko + i\, b\, \omega - m\, \omega^2\right)\right)\Big)\Big)\Big),$$
$$v \to (fp\, \kappa\, (\kappa\, \omega\, (b + i\, m\, \omega)\, (\sigma + i\, \alpha\, \omega) + 2\, ko\, \kappa\, (-i\, \sigma + \alpha\, \omega) + k\, (-i\, \gamma\, \kappa\, \sigma + \delta\, \omega - \gamma\, \delta\, \omega + \alpha\, \gamma\, \kappa\, \omega)))\Big/$$
$$\Big(2\, \Big(c\, k^2\, \gamma\, \kappa^2\, \omega + \omega\, (-i\, b + m\, \omega)\, \left(-2\, \delta^2\, \omega + \delta\, \kappa\, (-i\, \sigma + \alpha\, \omega) + c\, \kappa^2\, \omega \left(-2\, ko - i\, b\, \omega + m\, \omega^2\right)\right) -$$
$$i\, g\, \kappa^2\, \left(k^2\, \gamma + k\, (1 + \gamma)\, \left(ko + i\, b\, \omega - m\, \omega^2\right) + \omega\, (-i\, b + m\, \omega)\, \left(-2\, ko - i\, b\, \omega + m\, \omega^2\right)\right) +$$
$$k\, \Big(\omega\, \left(\delta^2 + c\, \kappa^2\, \left(ko + i\, b\, \omega - m\, \omega^2\right)\right) +$$
$$\gamma\, \left(\delta^2\, \omega + i\, \delta\, \kappa\, (\sigma + i\, \alpha\, \omega) + c\, \kappa^2\, \omega\, \left(ko + i\, b\, \omega - m\, \omega^2\right)\right)\Big)\Big)\Big)\Big\}\Big\}$$

In[∘]:=
```
u1 = -((ⅈ fp δ ω (2 δ^2 ω + c κ^2 ω (2 ko + k γ + ω (ⅈ b - m ω)) + g κ^2 (-2 ⅈ ko - ⅈ k γ + ω (b + ⅈ m ω)))) /
    (2 (δ (-ⅈ g ko κ^2 + (δ^2 + c ko κ^2) ω) (-k + ω (-ⅈ b + m ω)) - δ (k γ + ω (ⅈ b - m ω))
        (δ (ⅈ κ σ + δ ω - α κ ω) - ⅈ κ^2 (g + ⅈ c ω) (k + ko + ω (ⅈ b - m ω))))));
u2 = (fp ω (2 ⅈ δ^2 ω - 2 δ κ (σ + ⅈ α ω) +
      g κ^2 (k + 2 ko + ω (ⅈ b - m ω)) + ⅈ c κ^2 ω (k + 2 ko + ω (ⅈ b - m ω)))) /
   (2 (c k^2 γ κ^2 ω + ω (-ⅈ b + m ω) (-2 δ^2 ω + δ κ (-ⅈ σ + α ω) + c κ^2 ω (-2 ko - ⅈ b ω + m ω^2)) -
      ⅈ g κ^2 (k^2 γ + k (1 + γ) (ko + ⅈ b ω - m ω^2) + ω (-ⅈ b + m ω) (-2 ko - ⅈ b ω + m ω^2)) +
      k (ω (δ^2 + c κ^2 (ko + ⅈ b ω - m ω^2)) +
         γ (δ^2 ω + ⅈ δ κ (σ + ⅈ α ω) + c κ^2 ω (ko + ⅈ b ω - m ω^2)))));
v = (fp κ (κ ω (b + ⅈ m ω) (σ + ⅈ α ω) + 2 ko κ (-ⅈ σ + α ω) + k (-ⅈ γ κ σ + δ ω - γ δ ω + α γ κ ω))) /
   (2 (c k^2 γ κ^2 ω + ω (-ⅈ b + m ω) (-2 δ^2 ω + δ κ (-ⅈ σ + α ω) + c κ^2 ω (-2 ko - ⅈ b ω + m ω^2)) -
      ⅈ g κ^2 (k^2 γ + k (1 + γ) (ko + ⅈ b ω - m ω^2) + ω (-ⅈ b + m ω) (-2 ko - ⅈ b ω + m ω^2)) +
      k (ω (δ^2 + c κ^2 (ko + ⅈ b ω - m ω^2)) +
         γ (δ^2 ω + ⅈ δ κ (σ + ⅈ α ω) + c κ^2 ω (ko + ⅈ b ω - m ω^2)))));
Simplify[u2 / u1]
Simplify[v / u1]
```

Out[∘]=
$$\frac{2\,\delta^2\,\omega + 2\,i\,\delta\,\kappa\,(\sigma + i\,\alpha\,\omega) - i\,g\,\kappa^2\left(k + 2\,ko + i\,b\,\omega - m\,\omega^2\right) + c\,\kappa^2\,\omega\left(k + 2\,ko + i\,b\,\omega - m\,\omega^2\right)}{2\,\delta^2\,\omega + c\,\kappa^2\,\omega\,(2\,ko + k\,\gamma + \omega\,(i\,b - m\,\omega)) + g\,\kappa^2\,(-2\,i\,ko - i\,k\,\gamma + \omega\,(b + i\,m\,\omega))}$$

Out[∘]=
$$\frac{\kappa\,(2\,ko\,\kappa\,(\sigma + i\,\alpha\,\omega) + i\,\kappa\,\omega\,(b + i\,m\,\omega)\,(\sigma + i\,\alpha\,\omega) + k\,(i\,\delta\,\omega - i\,\gamma\,\delta\,\omega + \gamma\,\kappa\,(\sigma + i\,\alpha\,\omega)))}{\omega\left(-2\,\delta^2\,\omega + i\,g\,\kappa^2\,(2\,ko + k\,\gamma + \omega\,(i\,b - m\,\omega)) + c\,\kappa^2\,\omega\,(-2\,ko - k\,\gamma + \omega\,(-i\,b + m\,\omega))\right)}$$

In[∘]:=
```
ComplexExpand[(2 (δ (-ⅈ g ko κ^2 + (δ^2 + c ko κ^2) ω) (-k + ω (-ⅈ b + m ω)) -
      δ (k γ + ω (ⅈ b - m ω)) (δ (ⅈ κ σ + δ ω - α κ ω) - ⅈ κ^2 (g + ⅈ c ω) (k + ko + ω (ⅈ b - m ω)))))]
```

Out[∘]=
$$\begin{aligned}
&-2\,k\,\delta^3\,\omega - 2\,k\,\gamma\,\delta^3\,\omega + 2\,k\,\alpha\,\gamma\,\delta^2\,\kappa\,\omega - 2\,b\,g\,k\,\delta\,\kappa^2\,\omega - 4\,b\,g\,ko\,\delta\,\kappa^2\,\omega - 2\,c\,k\,ko\,\delta\,\kappa^2\,\omega - 2\,b\,g\,k\,\gamma\,\delta\,\kappa^2\,\omega - \\
&\quad 2\,c\,k^2\,\gamma\,\delta\,\kappa^2\,\omega - 2\,c\,k\,ko\,\gamma\,\delta\,\kappa^2\,\omega + 2\,b\,\delta^2\,\kappa\,\sigma\,\omega + 4\,m\,\delta^3\,\omega^3 - 2\,m\,\alpha\,\delta^2\,\kappa\,\omega^3 + 2\,b^2\,c\,\delta\,\kappa^2\,\omega^3 + \\
&\quad 4\,b\,g\,m\,\delta\,\kappa^2\,\omega^3 + 2\,c\,k\,m\,\delta\,\kappa^2\,\omega^3 + 4\,c\,ko\,m\,\delta\,\kappa^2\,\omega^3 + 2\,c\,k\,m\,\gamma\,\delta\,\kappa^2\,\omega^3 - 2\,c\,m^2\,\delta\,\kappa^2\,\omega^5 + \\
&\quad i\,\big(2\,g\,k\,ko\,\delta\,\kappa^2 + 2\,g\,k^2\,\gamma\,\delta\,\kappa^2 + 2\,g\,k\,ko\,\gamma\,\delta\,\kappa^2 - 2\,k\,\gamma\,\delta^2\,\kappa\,\sigma - 4\,b\,\delta^3\,\omega^2 + 2\,b\,\alpha\,\delta^2\,\kappa\,\omega^2 - \\
&\qquad 2\,b^2\,g\,\delta\,\kappa^2\,\omega^2 - 2\,b\,c\,k\,\delta\,\kappa^2\,\omega^2 - 4\,b\,c\,ko\,\delta\,\kappa^2\,\omega^2 - 2\,g\,k\,m\,\delta\,\kappa^2\,\omega^2 - 4\,g\,ko\,m\,\delta\,\kappa^2\,\omega^2 - \\
&\qquad 2\,b\,c\,k\,\gamma\,\delta\,\kappa^2\,\omega^2 - 2\,g\,k\,m\,\gamma\,\delta\,\kappa^2\,\omega^2 + 2\,m\,\delta^2\,\kappa\,\sigma\,\omega^2 + 4\,b\,c\,m\,\delta\,\kappa^2\,\omega^4 + 2\,g\,m^2\,\delta\,\kappa^2\,\omega^4\big)
\end{aligned}$$

In[∘]:=
```
Simplify[Solve[
  {-2 k δ^3 ω - 2 k γ δ^3 ω + 2 k α γ δ^2 κ ω - 2 b g k δ κ^2 ω - 4 b g ko δ κ^2 ω - 2 c k ko δ κ^2 ω - 2 b g k γ δ
      κ^2 ω - 2 c k^2 γ δ κ^2 ω - 2 c k ko γ δ κ^2 ω + 2 b δ^2 κ σ ω + 4 m δ^3 ω^3 - 2 m α δ^2 κ ω^3 + 2 b^2 c δ κ^2 ω^3 +
     4 b g m δ κ^2 ω^3 + 2 c k m δ κ^2 ω^3 + 4 c ko m δ κ^2 ω^3 + 2 c k m γ δ κ^2 ω^3 - 2 c m^2 δ κ^2 ω^5 == 0,
   (2 g k ko δ κ^2 + 2 g k^2 γ δ κ^2 + 2 g k ko γ δ κ^2 - 2 k γ δ^2 κ σ - 4 b δ^3 ω^2 + 2 b α δ^2 κ ω^2 -
      2 b^2 g δ κ^2 ω^2 - 2 b c k δ κ^2 ω^2 - 4 b c ko δ κ^2 ω^2 - 2 g k m δ κ^2 ω^2 - 4 g ko m δ κ^2 ω^2 - 2 b c k γ
       δ κ^2 ω^2 - 2 g k m γ δ κ^2 ω^2 + 2 m δ^2 κ σ ω^2 + 4 b c m δ κ^2 ω^4 + 2 g m^2 δ κ^2 ω^4) == 0}, {σ, α}]]
```

Out[∘]=
```
{{σ → (-b ω^2 (k (-1 + γ) δ^2 + c k^2 γ^2 κ^2 + c κ^2 ω^2 (b^2 + m^2 ω^2) + c k κ^2 (ko (-1 + γ) - 2 m γ ω^2)) +
      g κ^2 (k^3 γ^2 + ω^2 (2 ko - m ω^2) (b^2 + m^2 ω^2) + k^2 γ (ko (1 + γ) - m (2 + γ) ω^2) +
         k ω^2 (b^2 + m (-ko (1 + 3 γ) + m (1 + 2 γ) ω^2)))) / (δ κ (k^2 γ^2 + b^2 ω^2 - 2 k m γ ω^2 + m^2 ω^4)),
  α → (c k^3 γ^2 κ^2 + ω^2 (b^2 + m^2 ω^2) (2 δ^2 + κ^2 (b g + 2 c ko - c m ω^2)) +
      k^2 γ ((1 + γ) δ^2 + κ^2 (b g γ + c ko (1 + γ) - c m (2 + γ) ω^2)) +
      k (b^2 c κ^2 ω^2 + b g κ^2 (ko (-1 + γ) - 2 m γ ω^2) +
         m ω^2 (-((1 + 3 γ) δ^2) + c κ^2 (-ko (1 + 3 γ) + m (1 + 2 γ) ω^2)))) /
    (δ κ (k^2 γ^2 + b^2 ω^2 - 2 k m γ ω^2 + m^2 ω^4))}}
```

assymetric forcing case

In[∘]:=
```
ω =.;
b =.;
g =.;
σ =.;
α =.;
k1 = k;
k2 = k γ;
m1 = m;
m2 = m;

Simplify[Solve[{( (ko + k1) - m1 ω^2 + ⅈ b ω) u1/(I ω) - ko u2/(I ω) - v δ/κ == f_p,
   ( (ko + k2) - m2 ω^2 + ⅈ b ω) u2/(I ω) - ko u1/(I ω) + v δ/κ == 0,
   (I ω c + g) v + δ/κ (u1 - u2) == (α + σ/(I ω)) (u1)}, {u1, u2, v}]]
```

Out[∘]=
```
{{u1 → (ⅈ ω (ⅈ g δ κ^2 (ko + k γ + ⅈ b ω - m ω^2) - δ ω (δ^2 + c κ^2 (ko + k γ + ⅈ b ω - m ω^2))) f_p) /
    (δ (-ⅈ g ko κ^2 + (δ^2 + c ko κ^2) ω) (-k + ω (-ⅈ b + m ω)) -
      δ (k γ + ω (ⅈ b - m ω)) (δ (ⅈ κ σ + δ ω - α κ ω) - ⅈ κ^2 (g + ⅈ c ω) (k + ko + ω (ⅈ b - m ω)))),
  u2 → (ω (g ko κ^2 + ⅈ (δ^2 ω + c ko κ^2 ω + ⅈ δ κ (σ + ⅈ α ω))) f_p) /
    (c k^2 γ κ^2 ω + ω (-ⅈ b + m ω) (-2 δ^2 ω + δ κ (-ⅈ σ + α ω) + c κ^2 ω (-2 ko - ⅈ b ω + m ω^2)) -
      ⅈ g κ^2 (k^2 γ + k (1 + γ) (ko + ⅈ b ω - m ω^2) + ω (-ⅈ b + m ω) (-2 ko - ⅈ b ω + m ω^2)) +
      k (ω (δ^2 + c κ^2 (ko + ⅈ b ω - m ω^2)) + γ (δ^2 ω + ⅈ δ κ (σ + ⅈ α ω) + c κ^2 ω (ko + ⅈ b ω - m ω^2)))),
  v → (κ (ko κ (-ⅈ σ + α ω) + (ⅈ κ σ + δ ω - α κ ω) (-k γ + ω (-ⅈ b + m ω))) f_p) /
    (c k^2 γ κ^2 ω + ω (-ⅈ b + m ω) (-2 δ^2 ω + δ κ (-ⅈ σ + α ω) + c κ^2 ω (-2 ko - ⅈ b ω + m ω^2)) -
      ⅈ g κ^2 (k^2 γ + k (1 + γ) (ko + ⅈ b ω - m ω^2) + ω (-ⅈ b + m ω) (-2 ko - ⅈ b ω + m ω^2)) + k
       (ω (δ^2 + c κ^2 (ko + ⅈ b ω - m ω^2)) + γ (δ^2 ω + ⅈ δ κ (σ + ⅈ α ω) + c κ^2 ω (ko + ⅈ b ω - m ω^2))))}}
```

check denominator equivalence -- yes

```
Simplify[((δ (-i g ko κ^2 + (δ^2 + c ko κ^2) ω) (-k + ω (-i b + m ω)) -
      δ (k γ + ω (i b - m ω)) (δ (i κ σ + δ ω - α κ ω) - i κ^2 (g + i c ω) (k + ko + ω (i b - m ω))))) /
  (δ (-i g ko κ^2 + (δ^2 + c ko κ^2) ω) (-k + ω (-i b + m ω)) -
     δ (k γ + ω (i b - m ω)) (δ (i κ σ + δ ω - α κ ω) - i κ^2 (g + i c ω) (k + ko + ω (i b - m ω))))]
Simplify[((c k^2 γ κ^2 ω + ω (-i b + m ω) (-2 δ^2 ω + δ κ (-i σ + α ω) + c κ^2 ω (-2 ko - i b ω + m ω^2)) -
      i g κ^2 (k^2 γ + k (1 + γ) (ko + i b ω - m ω^2) + ω (-i b + m ω) (-2 ko - i b ω + m ω^2)) +
      k (ω (δ^2 + c κ^2 (ko + i b ω - m ω^2)) + γ (δ^2 ω + i δ κ (σ + i α ω) + c κ^2 ω (ko + i b ω - m ω^2))))) /
  (c k^2 γ κ^2 ω + ω (-i b + m ω) (-2 δ^2 ω + δ κ (-i σ + α ω) + c κ^2 ω (-2 ko - i b ω + m ω^2)) -
     i g κ^2 (k^2 γ + k (1 + γ) (ko + i b ω - m ω^2) + ω (-i b + m ω) (-2 ko - i b ω + m ω^2)) +
     k (ω (δ^2 + c κ^2 (ko + i b ω - m ω^2)) + γ (δ^2 ω + i δ κ (σ + i α ω) + c κ^2 ω (ko + i b ω - m ω^2))))]
```

Out[∘]= 1

Out[∘]= 2

```
In[∘]:= ComplexExpand[(δ (k + ω (i b - m ω)) (δ^2 ω + κ^2 (-i g + c ω) ko) +
     δ (k γ + ω (i b - m ω)) (δ (i κ σ + δ ω - α κ ω) - i κ^2 (g + i c ω) (k + ω (i b - m ω) + ko)))]
```

```
Out[∘]= k δ^3 ω + k γ δ^3 ω - k α γ δ^2 κ ω + b g k δ κ^2 ω + 2 b g ko δ κ^2 ω + c k ko δ κ^2 ω +
  b g k γ δ κ^2 ω + c k^2 γ δ κ^2 ω + c k ko γ δ κ^2 ω - b δ^2 κ σ ω - 2 m δ^3 ω^3 + m α δ^2 κ ω^3 -
  b^2 c δ κ^2 ω^3 - 2 b g m δ κ^2 ω^3 - c k m δ κ^2 ω^3 - 2 c ko m δ κ^2 ω^3 - c k m γ δ κ^2 ω^3 + c m^2 δ κ^2 ω^5 +
  i (-g k ko δ κ^2 - g k^2 γ δ κ^2 - g k ko γ δ κ^2 + k γ δ^2 κ σ + 2 b δ^3 ω^2 - b α δ^2 κ ω^2 +
     b^2 g δ κ^2 ω^2 + b c k δ κ^2 ω^2 + 2 b c ko δ κ^2 ω^2 + g k m δ κ^2 ω^2 + 2 g ko m δ κ^2 ω^2 +
     b c k γ δ κ^2 ω^2 + g k m γ δ κ^2 ω^2 - m δ^2 κ σ ω^2 - 2 b c m δ κ^2 ω^4 - g m^2 δ κ^2 ω^4)
```

```
In[∘]:= ω =.;
Simplify[Solve[{k δ^3 ω + k γ δ^3 ω - k α γ δ^2 κ ω + b g k δ κ^2 ω + 2 b g ko δ κ^2 ω + c k ko δ κ^2 ω +
       b g k γ δ κ^2 ω + c k^2 γ δ κ^2 ω + c k ko γ δ κ^2 ω - b δ^2 κ σ ω - 2 m δ^3 ω^3 + m α δ^2 κ ω^3 - b^2 c δ κ^2 ω^3 -
       2 b g m δ κ^2 ω^3 - c k m δ κ^2 ω^3 - 2 c ko m δ κ^2 ω^3 - c k m γ δ κ^2 ω^3 + c m^2 δ κ^2 ω^5 == 0,
     (-g k ko δ κ^2 - g k^2 γ δ κ^2 - g k ko γ δ κ^2 + k γ δ^2 κ σ + 2 b δ^3 ω^2 - b α δ^2 κ ω^2 + b^2 g δ κ^2 ω^2 +
        b c k δ κ^2 ω^2 + 2 b c ko δ κ^2 ω^2 + g k m δ κ^2 ω^2 + 2 g ko m δ κ^2 ω^2 + b c k γ δ κ^2 ω^2 +
        g k m γ δ κ^2 ω^2 - m δ^2 κ σ ω^2 - 2 b c m δ κ^2 ω^4 - g m^2 δ κ^2 ω^4) == 0}, {σ, α}]]
```

```
Out[∘]= {{σ → (-b ω^2 (k (-1 + γ) δ^2 + c k^2 γ^2 κ^2 + c κ^2 ω^2 (b^2 + m^2 ω^2) + c k κ^2 (ko (-1 + γ) - 2 m γ ω^2)) +
        g κ^2 (k^3 γ^2 + ω^2 (2 ko - m ω^2) (b^2 + m^2 ω^2) + k^2 γ (ko (1 + γ) - m (2 + γ) ω^2) +
           k ω^2 (b^2 + m (-ko (1 + 3 γ) + m (1 + 2 γ) ω^2)))) / (δ κ (k^2 γ^2 + b^2 ω^2 - 2 k m γ ω^2 + m^2 ω^4)),
   α → (c k^3 γ^2 κ^2 + ω^2 (b^2 + m^2 ω^2) (2 δ^2 + κ^2 (b g + 2 c ko - c m ω^2)) +
        k^2 γ ((1 + γ) δ^2 + κ^2 (b g γ + c ko (1 + γ) - c m (2 + γ) ω^2)) +
        k (b^2 c κ^2 ω^2 + b g κ^2 (ko (-1 + γ) - 2 m γ ω^2) +
           m ω^2 (-((1 + 3 γ) δ^2) + c κ^2 (-ko (1 + 3 γ) + m (1 + 2 γ) ω^2)))) /
     (δ κ (k^2 γ^2 + b^2 ω^2 - 2 k m γ ω^2 + m^2 ω^4))}}
```

Confirm symmetric and asymmetric cases give same alpha and sigma -- yes

```
In[∘]:= Simplify[(-b ω^2 (k (-1 + γ) δ^2 + c k^2 γ^2 κ^2 + c κ^2 ω^2 (b^2 + m^2 ω^2) + c k κ^2 (ko (-1 + γ) - 2 m γ ω^2)) +
          g κ^2 (k^3 γ^2 + ω^2 (2 ko - m ω^2) (b^2 + m^2 ω^2) + k^2 γ (ko (1 + γ) - m (2 + γ) ω^2) +
             k ω^2 (b^2 + m (-ko (1 + 3 γ) + m (1 + 2 γ) ω^2)))) / (δ κ (k^2 γ^2 + b^2 ω^2 - 2 k m γ ω^2 + m^2 ω^4)) -
      (-b ω^2 (k (-1 + γ) δ^2 + c k^2 γ^2 κ^2 + c κ^2 ω^2 (b^2 + m^2 ω^2) + c k κ^2 (ko (-1 + γ) - 2 m γ ω^2)) +
          g κ^2 (k^3 γ^2 + ω^2 (2 ko - m ω^2) (b^2 + m^2 ω^2) + k^2 γ (ko (1 + γ) - m (2 + γ) ω^2) +
             k ω^2 (b^2 + m (-ko (1 + 3 γ) + m (1 + 2 γ) ω^2)))) / (δ κ (k^2 γ^2 + b^2 ω^2 - 2 k m γ ω^2 + m^2 ω^4))]

Out[∘]= 0

In[∘]:= Simplify[(c k^3 γ^2 κ^2 + ω^2 (b^2 + m^2 ω^2) (2 δ^2 + κ^2 (b g + 2 c ko - c m ω^2)) +
          k^2 γ ((1 + γ) δ^2 + κ^2 (b g γ + c ko (1 + γ) - c m (2 + γ) ω^2)) +
          k (b^2 c κ^2 ω^2 + b g κ^2 (ko (-1 + γ) - 2 m γ ω^2) +
             m ω^2 (-((1 + 3 γ) δ^2) + c κ^2 (-ko (1 + 3 γ) + m (1 + 2 γ) ω^2)))) /
      (δ κ (k^2 γ^2 + b^2 ω^2 - 2 k m γ ω^2 + m^2 ω^4)) - (c k^3 γ^2 κ^2 +
          ω^2 (b^2 + m^2 ω^2) (2 δ^2 + κ^2 (b g + 2 c ko - c m ω^2)) +
          k^2 γ ((1 + γ) δ^2 + κ^2 (b g γ + c ko (1 + γ) - c m (2 + γ) ω^2)) +
          k (b^2 c κ^2 ω^2 + b g κ^2 (ko (-1 + γ) - 2 m γ ω^2) + m ω^2 (-((1 + 3 γ) δ^2) +
                c κ^2 (-ko (1 + 3 γ) + m (1 + 2 γ) ω^2)))) / (δ κ (k^2 γ^2 + b^2 ω^2 - 2 k m γ ω^2 + m^2 ω^4))]

Out[∘]= 0
```
